# From the formation of embryonic appendages to the color of wings: Conserved and novel roles of *aristaless1* in butterfly development

**DOI:** 10.1101/2021.12.02.470931

**Authors:** Erick X. Bayala, Nicholas VanKuren, Darli Massardo, Marcus Kronforst

## Abstract

Highly diverse butterfly wing patterns have emerged as a powerful system for understanding the genetic basis of phenotypic variation. While the genetic basis of this pattern variation is being clarified, the precise developmental pathways linking genotype to phenotype are not well understood. The gene *aristaless*, which plays a role in appendage patterning and extension, has been duplicated in Lepidoptera. One copy, *aristaless1*, has been shown to control a white/yellow color switch in the butterfly *Heliconius cydno*, suggesting a novel function associated with color patterning and pigmentation. Here we investigate the developmental basis of *al1* in embryos, larvae and pupae using new antibodies, CRISPR/Cas9, RNAi, qPCR assays of downstream targets and pharmacological manipulation of an upstream activator. We find that Al1 is expressed at the distal tips of developing embryonic appendages consistent with its ancestral role. In developing wings, we observe Al1 accumulation within developing scale cells of white *H. cydno* during early pupation while yellow scale cells exhibit little Al1 at this timepoint. Reduced Al1 expression is also associated with yellow scale development in *al1* knockouts and knockdowns. We also find that Al1 expression appears to downregulate the enzyme Cinnabar and other genes that synthesize and transport the yellow pigment, 3–Hydroxykynurenine (3-OHK). Finally, we provide evidence that Al1 activation is under the control of Wnt signaling. We propose a model in which high levels of Al1 during early pupation, which are mediated by Wnt, are important for melanic pigmentation and specifying white portions of the wing while reduced levels of Al1 during early pupation promote upregulation of proteins needed to move and synthesize 3-OHK, promoting yellow pigmentation. In addition, we discuss how the ancestral role of *aristaless* in appendage extension may be relevant in understanding the cellular mechanism behind color patterning in the context of the heterochrony hypothesis.

## Introduction

The diversity and complexity of butterfly color patterns is striking. What is even more impressive is that this color pattern diversity within butterflies is often controlled by a small number of genes (Deshmukh, et al., 2017). Despite the importance of these color patterning genes for the life history and ecology of butterflies, we know very little about how similar or different these genes function during wing color pattern development. *Heliconius* butterflies are a great system to address this issue. In this genus, a handful of genes control the evolution and diversity of multiple color patterns (Kronforst & Papa, 2015; Van Belleghem, et al., 2017). One example is the signaling ligand *wntA*, which is expressed early within the larval wing imaginal discs and specifies future black patterns on the adult wing (Martin, et al., 2012, **Figure 1**). Another example is the transcription factor *optix*, which controls red color patterns across *Heliconius* by localizing within the nucleus of scale building cells during mid pupation (Reed, et al., 2011; Martin, et al., 2014, **Figure 1**). One last example is the gene *cortex,* which is a cell cycle regulator involved in the specification of melanic elements of the wing (Nadeau, et al., 2016). Despite major developmental differences and although cortex knock-outs may have more widespread effects on scale development (Livraghi, et al., 2021), all three of these genes have expression patterns that spatially prefigure future adult black and red color pattern elements at different stages of wing development. In addition to black and red patterns, multiple *Heliconius* species vary in color on light portions of their wings, specifically whether these scales are white (unpigmented) or yellow (containing the hemolymph derived pigment 3-hydroxykynurenine [3-OHK]; Gilbert, et al., 1988). Recently, the genetic switch between white and yellow scale fates in *Heliconius cydno*, which has historically been referred to as the *K* locus (Kronforst, et al., 2006; Chamberlain, et al., 2009), was traced back to the gene *aristaless1* (*al1*) in *Heliconius cydno* (Westerman, et al., 2018). However, we know little about the developmental basis of *al1* color switching, including how and when during development this gene controls the decision between white and yellow color phenotypes. Furthermore, we have no information about how the developmental biology of *al1* compares to *optix*, *wntA,* and *cortex* and if any general developmental trends, like the spatial prefiguring often described for these other genes, will emerge in the context of *Heliconius* color patterning.

**Figure 1:**
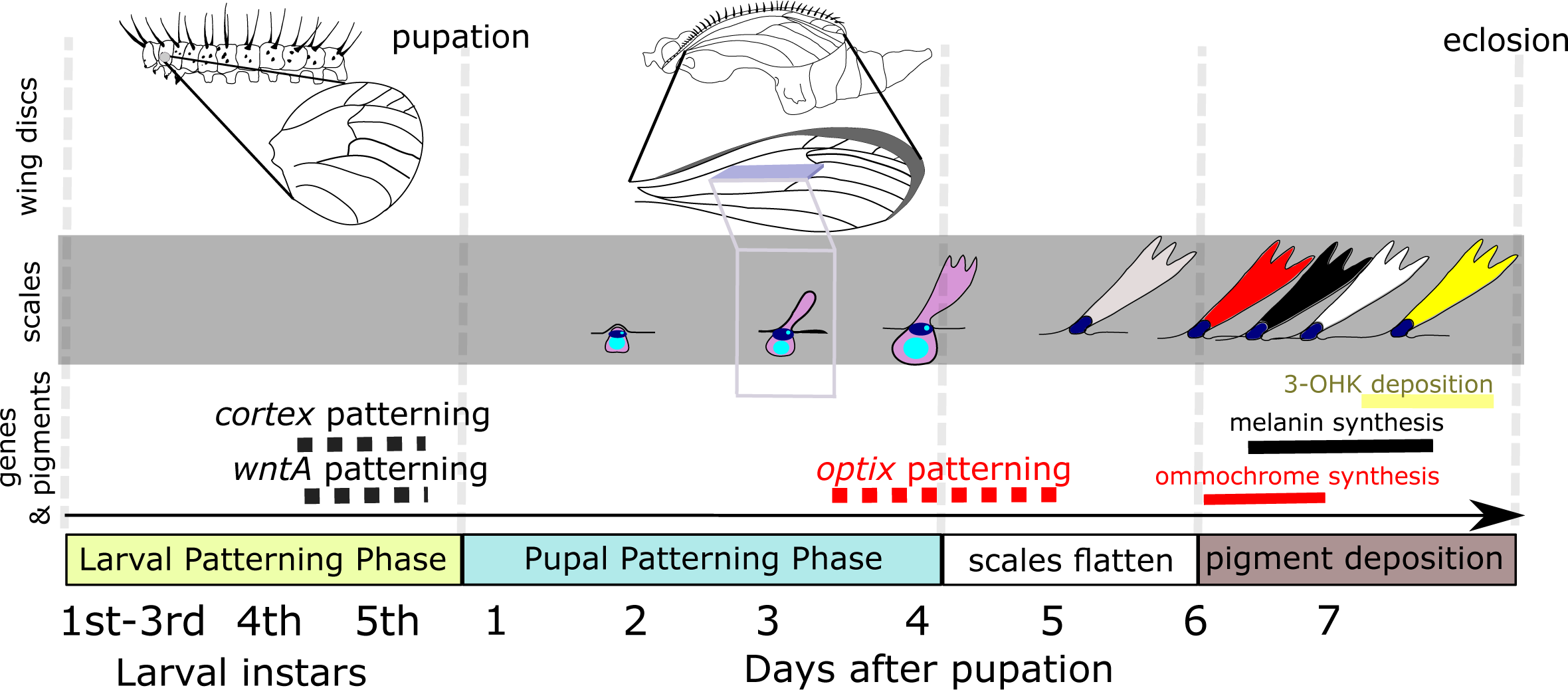
Summary of *Heliconius* wing pattern development. The top panel highlights the wing imaginal discs across the multiple phases of wing development at the organismal level. The middle panel describes developmental changes observed in the functional cells (scale cell in magenta and socket cell in dark blue) that will eventually become the pigmented scales (stages of scale development adapted from Dinwiddie, et al., 2014). The bottom panel of the image consolidates our knowledge about when the known patterning genes *wntA*, *cortex* and *optix* are expressed and when the expression results in terminal color synthesis of melanin and ommochromes, respectively. The yellow pigment (3-OHK) deposition window is also shown. Dashed gray lines separete the different phases.

Here we investigate how *al1* specifies white and yellow wing coloration by studying the timing of *al1* transcription and protein localization in developing wings of the butterfly *Heliconius cydno*, a species with polymorphic wing coloration. The homeobox transcription factor *aristaless1* is one of two paralogs stemming from a gene duplication event that occurred at the base of Lepidoptera (Martin and Reed, 2010). Much of what we know about the single-copy ancestral *aristaless* (*al*) comes from work in *Drosophila* and shows that it is often associated with the extension and patterning of appendages. (Schneitz, et al., 1993). Gene expression studies in flies (Campbell & Tomlinson, 1988; Schneitz, et al., 1993) have shown that *al* accumulates along the distal edges of extending structures such as leg, wing, and antennae during different developmental stages. Furthermore, knockouts of *al* in flies (Schneitz, et al., 1993) often result in malformed or missing distal elements of appendages. These observations in *Drosophila* have been reinforced in other insects like beetles (Moczek, 2005) and crickets (Beermann and Schroder, 2004; Miyawaki, et al., 2002). There is also some information on the developmental role of *al1* in Lepidoptera. For instance, in the moth *Bombyx mori, al1* has been shown to be crucial for the extension and branching patterns of antennae (Ando, et al., 2018). In this example, *al1* expression and protein localization were observed within all of the extending branches of the antennae (Ando, et al 2018). In addition, in some nymphalid butterflies *al2* has been shown to play a role in specifying melanic discal (black patches in the middle of the wing) color pattern elements on the wing (Martin and Reed, 2010). In summary, *al* has been described on multiple occasions and across several organisms as a key regulator of developmental processes. Previous descriptions of *al1*’s role in the extension of appendages and perhaps wing patterning beg the question of how this gene mediates the developmental decision between white and yellow wing patterns in *Heliconius* butterflies.

Here we analyze CRISPR/Cas9 knockouts in adult wings to describe the multiple effects that *al1* has on color patterning in *Heliconius*. We also use a combination of staining techniques to describe Al1 subcellular localization first in embryos appendages, and then across the development of the wing in order to determine when and where Al1 may be controlling the decision between white and yellow color patterns. Then, we combine knockout and knockdown approaches with our Al1 staining to provide functional evidence for how Al1 subcellular localization relates to the final specification of color pattern. Finally, we perform quantitative PCR (qPCR) analyses to determine possible downstream genes under the control of Al1 and employ a pharmacological agent to dissect the role of an upstream pathway in the regulation of Al1. Our results reveal how *al1* controls white and yellow color patterns formation (specification to pigmentation) in *Heliconius* and help explain the developmental mechanisms leading to a fully pigmented *Heliconius* wing.

## Results

### *al1* knockouts switch white scales to yellow and black scales to brown but have no effect on yellow scales

Previous work used CRISPR/Cas9 knockouts to functionally test the involvement of *al1* in the switch between white and yellow wing color in *Heliconius cydno* (Westerman, et al., 2018). In these experiments, genetically white *H. cydno* with an *al1* knockout exhibited a switch of white scales to yellow scales (Westerman, et al., 2018).

To study the developmental role of *al1* we generated new CRISPR/Cas9 knockouts and recovered both the previously described as well as novel effects. As previously described, *al1* knockout clones within the white band of a genetically white *H. cydno* switched white scales to yellow (**Figure 2A**). However, careful observation of these yellow clones in white *H. cydno* revealed that when these clones expanded over the melanic regions of the wing, black scales became brown (**Figure 2B**). Previous work reported that Al1 seemed to be acting as a repressor of the yellow fate (Westerman, et al., 2018). Based on this repressor activity we hypothesized that *al1* knockout clones in genetically yellow *H. cydno* would have no effect on the yellow portions of the wing. In favor of this hypothesis we did not see any effects on the yellow parts of the wing, yet interestingly, similar to white butterflies, clones within the melanic regions of yellow butterflies also exhibited a switch from black to brown scales (**Figure 2B**).

**Figure 2:**
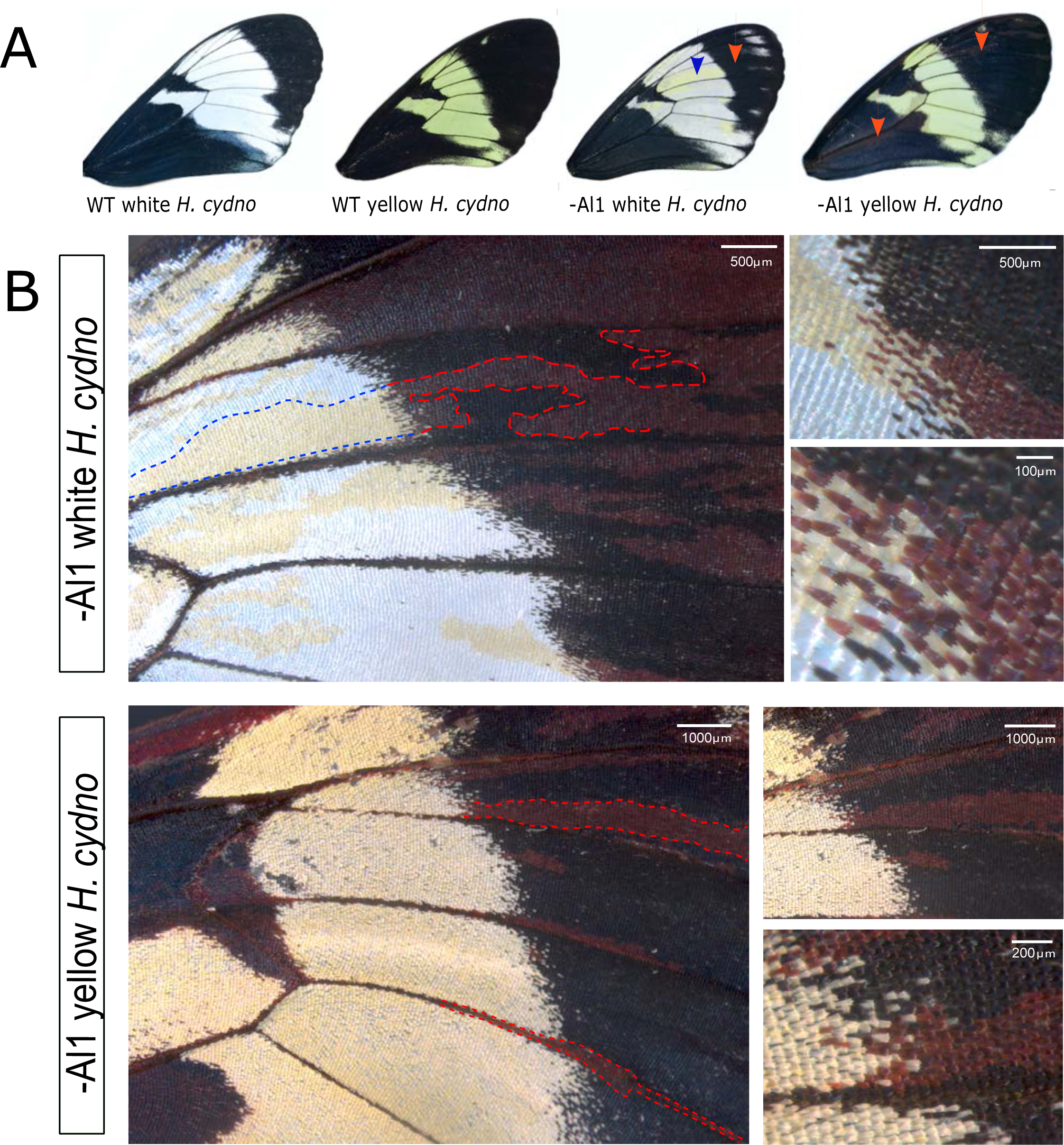
Wild type and *al1* CRISPR/Cas9 knockout forewings of white and yellow *H. cydno*. (**A**) Full adult forewing view of wild type and *al1* knockouts of both white and yellow *H. cydno*. Blue arrowheads highlight mutant yellow clones inside the white regions and red arrowheads highlight mutant brown clones inside the melanic regions of wing. (**B**) Higher magnification view of the mutant parts of the wing for both white (top panel) and yellow (bottom panel) *H. cydno* butterflies. Dashed blue lines highlight the parts of the clone that switched from white to yellow and dashed red lines highlight the parts of a clone that switched from black to brown.

These results confirm the importance of *al1* for the development of white wing coloration. If *al1* is knocked out, scales then switch to the yellow fate. However, the newly described *al1* knockout effects in melanic regions suggest a general role of *al1* in scale development across the entire wing, not just in the white/yellow band. Based on the widespread effect observed in white *H. cydno,* we hypothesized that *al1* expression may be important for scale development across the entire wing except for the yellow band of yellow *H. cydno*. We tested this hypothesis by analyzing *al1* expression and protein localization across multiple developmental stages for both yellow and white *H. cydno* butterflies.

### Al1 staining in embryos recapitulates the previous known role of Al with respect to proper appendage extension

Most of the previous Al1 work in nymphalid butterflies was done using the DP311 antibody, which is known to stain homeodomain transcription factors like Al1. However, this reagent is known to cross-react with similar proteins like the paralog Aristaless2 (Martin and Reed, 2010). In order to avoid this, we developed specific antibodies against *H. cydno* Al1 epitopes to determine the protein subcellular localization and pattern of expression in wings (**Figure S2**).

Before looking into Al1 expression pattern in wings, we tested our antibody specificity in *Heliconius cydno* embryos where we analyzed its relationship relative to the ancestral Al function in appendages. We also aimed to provide expectations of its subcellular localization within appendages as a point of comparison for wings. Similar to what has been reported in other insect systems (Campbell & Tomlinson, 1988; Schneitz, et al., 1993; Miyawaki, et al., 2002; Beermann and Schroder, 2004) for Al, we observed Al1 localized on the distal tip of appendages extending out of the primary body plan (**Figure 3A**). We observed accumulation within the cellular buds giving rise to the mouthparts within the head region (**Figure 3B**). In addition, we observed a clear accumulation of Al1 within the distal tips of the thoracic (**Figure 3C**), abdominal (**Figure 3D**), and anal prolegs. We also observed accumulation on the dorsal side of the embryo which has not previously been described in other systems. Surprisingly higher magnification revealed no apparent co-localization with the nucleus of cells at the distal tips (**Figure 3B-D**). To further elucidate our antibody specificity and determine if Al1 expression was causally related to appendage extension, we stained CRISPR Al1 knockout embryos. We observed sections of the embryos depleted for Al1, as expected from a CRISPR knockout (**Figure 3E-G**). In addition, areas depleted of Al1 exhibited elongation defects when compared to the same appendages within the embryos that had normal levels of Al1. In addition to confirming a role for Al1 in appendage extension in *Heliconius* embryos, these data also provide evidence for the specificity of our newly developed antibodies, allowing us to further probe the role of Al1 in wing color patterning

**Figure 3:**
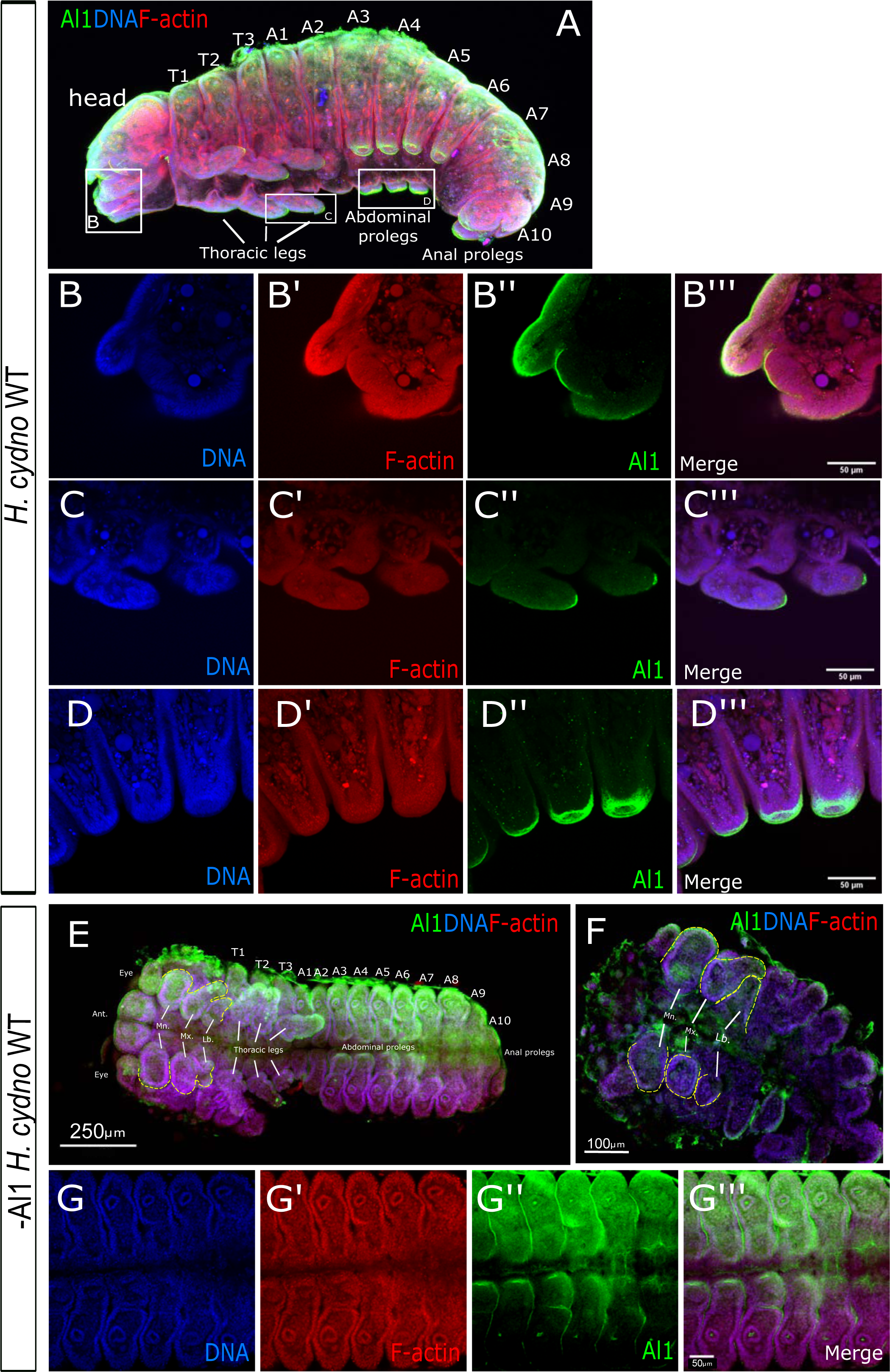
Immunodetection of Aristaless1 in wild-type and -Al1 CRISPR *Heliconius cydno* embyos. (**A**) Immunodetection of Al1 in wild-type embryos. White boxes highlight the mandibula (**B**), thoracic legs (**C**), and abdominal legs (**D**) zones shown at a higher magnification in the next panels. (**E-F**) Immunodetection of Al1 in injected -Al1 CRISPR embryos. (**G**) Higher magnification of the abdominal prolegs showcasing a zone lacking Al1. The segments and appendages are labeled for the full view embryos (**A, E-F**). Full embryo views highlight Antennal (Ant.), eyes, Mandibular (Mn), Maxillar (Mx), and Labial (Lb) head appendage precursors. The 3 pairs of thoracic legs, 4 pairs of abdominal prolegs, and the pair of anal prolegs buds are also marked. Panels show detection of DNA (B,C,D,G), F-actin (B’,C’,D’,G’), Al1 (B’’,C’’, D’, G’’), and a merge (A, B’’’,C’’,D’’,E-F,G’’’).

### Al1 accumulates in future white and black scale cell precursors, but not yellow scale cell precursors

Previous work with other nymphalid butterflies has shown that *al1* expression on larval wing discs resembles a modified pattern of the *aristaless* gene in flies (Martin & Reed, 2010). Using *in situ* hybridization and antibody staining, we found a similar pattern of expression of *al1* during larval wing disc development in white and yellow *H. cydno* (**Figure S1**). This expression pattern appears to be unrelated to the white vs. yellow color decision, hence we switched our attention to pupal stages.

Based on our CRISPR/Cas9 results, we hypothesized that Al1 would be present more widely across the wing, including the forewing band, of white *H. cydno* but would be absent from the band in yellow *H. cydno.* Furthermore, quantitative real-time PCR suggested that *al1* is expressed at all pupal stages but generally increases over time (Westerman et al., 2018). We therefore analyzed wings ranging from 2 days to 4 days (before scales harden and become impermeable to antibodies, **Figure 1**) after pupal formation (APF). We aimed our dissections to the 3 days APF mark because it allowed an efficient dissection without compromising the integrity of the wing and staining before any impermeability happens. In white *H. cydno* imaginal discs (Day 3 APF), Al1 was localized in developing scale cells for both future white and black scales (**Figure 4A-D**).

**Figure 4:**
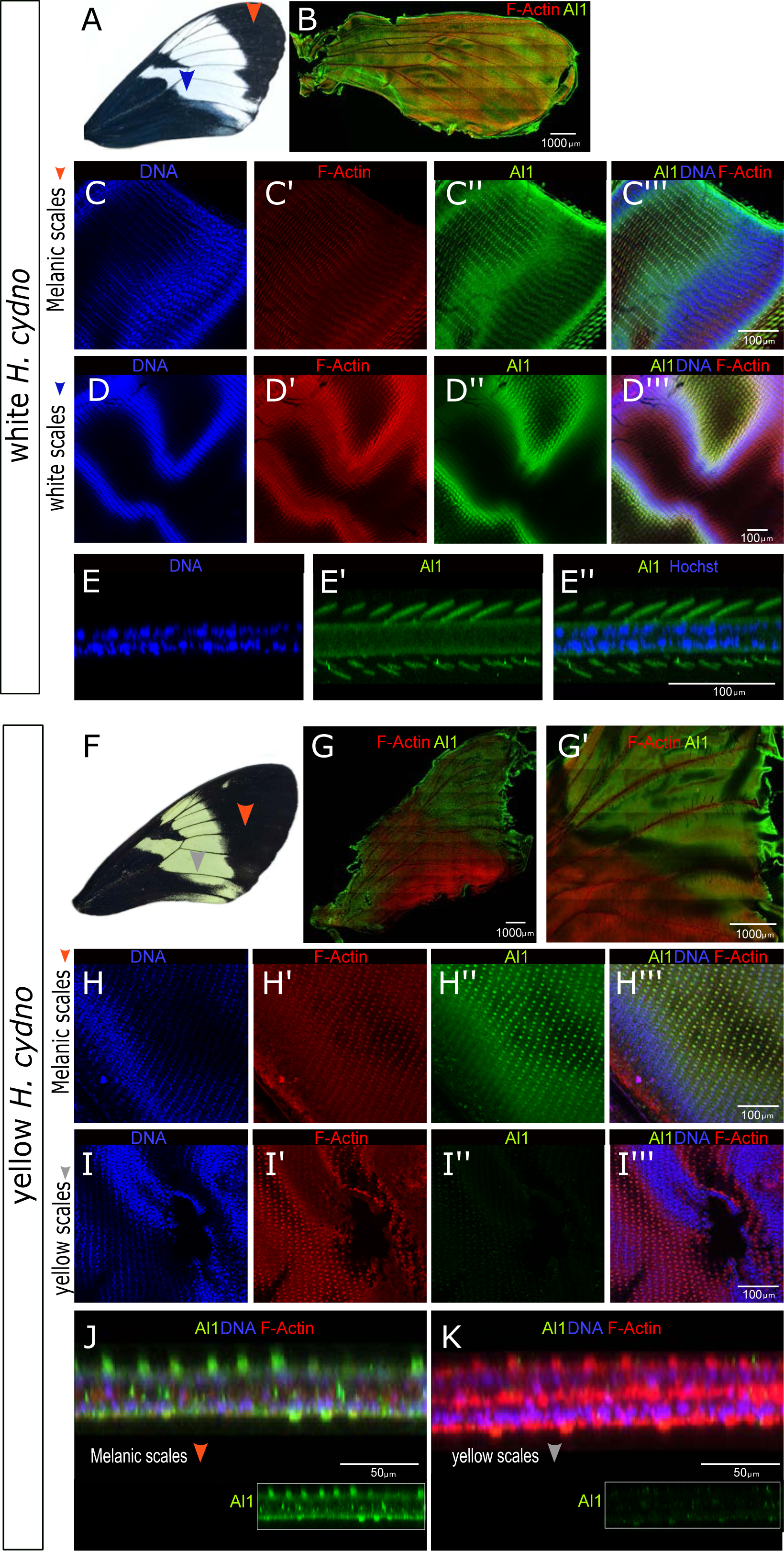
Immunodetection of Aristaless1 in white and yellow *Heliconius cydno* pupal forewings. (**A**) Adult forewing of a white *Heliconius cydno*. (**B**) Al1 detection in a full pupal wing of a white *Heliconius cydno* (3 days APF). (**C**) Details of Al1, DNA and actin detection in precursor scale cells of future melanic scales from a white *Heliconius cydno*. (**D**) Al1 detection in precursor scale cells of future white scales. (**E**) Side digital reconstruction from z-stack showing Al1 within precursor scale cells from the white part of the wing. (**F**) Adult forewing of a yellow *Heliconius cydno*. (**G**) Al1 detection in a full pupal wing of a yellow *Heliconius cydno*. **H**) Details of Al1 detection in precursor scale cells of future melanic scales from a yellow *Heliconius cydno*. (**I**) Al1 detection in precursor scale cells of future yellows scales. (**J-K**) Side digital reconstruction from z-stack showing differences in Al1 detection within precursor scale cells from yellow and melanic portions of a yellow *Heliconius cydno* wing. Panels show detection of DNA (C,D,E,H,I), F-actin (C’,D’,H’,I’), Al1 (C’’,D’’, E’, H’’,I’’), F-actin/DNA (B, G) and a merge (C’’’, D’’’,E’’,H’’,I’’’,J-K) view. Scale bars: B, G are 1000 μm; C-E and H-I are 100 μm; J-K are 50 μm.

This localization of Al1 was observed everywhere across the pupal wing on both the dorsal and ventral sides. Al1 did not appear to co-localize with the scale cell nucleus when analyzing multiple vertical planes (**Figure 4A-D**) similar to what we observed in embryo appendages (**Figure 3**). Careful observation of a side reconstruction from Z-stacks highlights that Al1 was concentrated within the cytoplasm of scale cells and absent, at least during these time-points, within the nucleus (**Figure 4E**). In contrast, Al1 was reduced or absent inside developing yellow scales (**Figure 4F-K**). This lack or lower levels of Al1 was more apparent during younger time points (day 2 to early day 3) and restricted to the dorsal side of the wing (**Figure S3**). Furthermore, as development continued, the overall level of Al1 on the dorsal side of yellow wings faded relative to that on the ventral side and this was not observed on white *H. cydno* wings (**Figure S3**). Using the vein patterns we inferred boundaries between future yellow and melanic parts of the wing and found a decrease in fluorescence associated with the transition from the melanic part of the wing to the yellow band (**Figure 5**).

**Figure 5:**
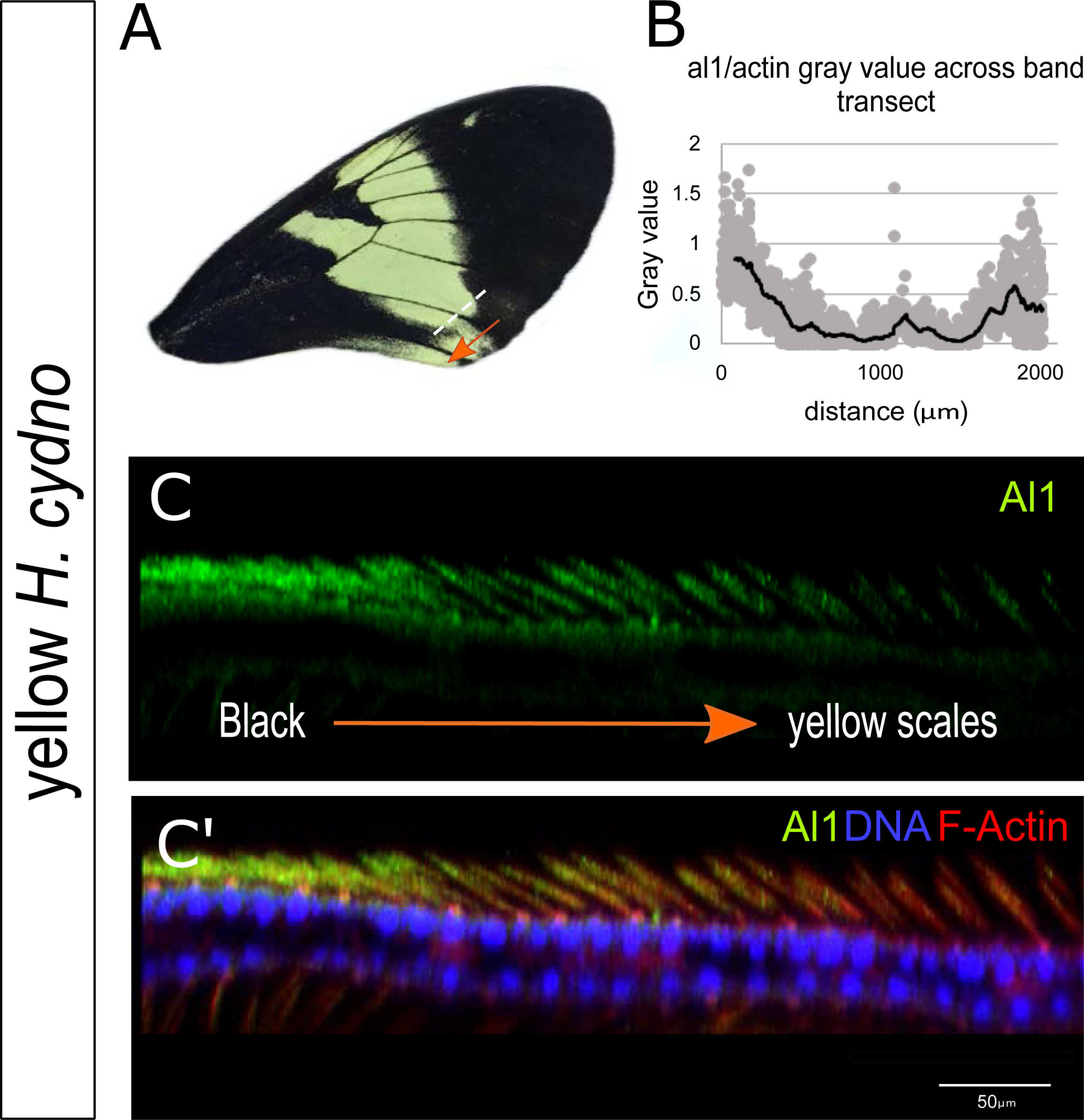
Immunodetection of Aristaless1 at the boundary between black and yellow scales in *Heliconius cydno* pupal forewing imaginal discs (3 Days APF). (**A**) Dorsal view of an adult yellow *H. cydno* forewing. (**B**) Quantification of the pixel gray value of a transect spanning across the presumptive yellow patch flanked by melanic regions at the stage of 3 Days APF. (**C**) Side view digital reconstruction from z-stack to observe the Al1 detection at the boundary between future melanic and yellow scales. Panel show detection Al1 (**C**) and a merged version (***C’***) with DNA and F-Actin detection. The orange arrow indicates the adult corresponding orientation for both the transect (white dashed line) for the B panel and the Z-stack of the side reconstruction of C. Scale bars: C, 50 μm.

Al1 is a homeodomain transcription factor and so we tested if it co-localized with the nucleus of scale cells at a later time point. Specifically, we examined wings at 4 days APF. In contrast, we found that white and black scales in white *H. cydno* again showed high levels Al1 in the cytoplasm of scale cells but not in the nucleus (**Figure S4**). Similarly, yellow *H. cydno* wings did not show nuclear localization of Al1 in the future melanic regions either (**Figure S4)**. We found no evidence that Al1 ever localized to the nucleus at 2 to 4 days APF, yet it is still possible that nuclear localization does occur at a time point that we did not observe or were not able to analyze. We verified antibody specificity by performing several negative controls and repeating staining in white *H. cydno* butterflies with antibodies against two different Al1 epitopes (**Figure S5A-D**).

These results suggest that the presence of Al1 in scale cells may be relevant for scale development and pigmentation across the entire wing. Presence of Al1 in the non-melanic band (which has already been specified by other genes like *wntA* [Martin, et al., 2012]) inhibits pigmentation resulting in white scales while absence or lower levels of Al1 in these developing scales during a short window early in pupation results in the switch of white scales to yellow scales. To further test this hypothesis, we examined Al1 expression in CRISPR/Cas9 knockouts and RNA interference (RNAi) knockdowns, which allowed us to directly correlate changes in protein localization with adult phenotype.

### Al1 CRISPR knockouts and RNA interference knockdowns reduce levels of Al1 and recapitulate the white to yellow color switch

To test our hypothesis that reduced or absent Al1 promote the switch from white to yellow, we determined Al1 levels by antibody staining in white *H. cydno* pupal wings with *al1* CRISPR/Cas9 knockouts (70% of the adult wings showed some level of mosaic color switch phenotype). Pupal wings analyzed at 3 days APF exhibited a depletion of Al1 in patches across the wing (**Figure 6**). Our observations with adult butterflies suggest that these clones lacking Al1 result in the switch of white and black scales to yellow and brown, respectively. We also characterized the range of CRISPR clone size and shape by observing a large number of CRISPR clones across the wings of white H. cydno, both in adults (**Figure S6**) and by antibody staining pupal wings (**Figure S7**).

**Figure 6:**
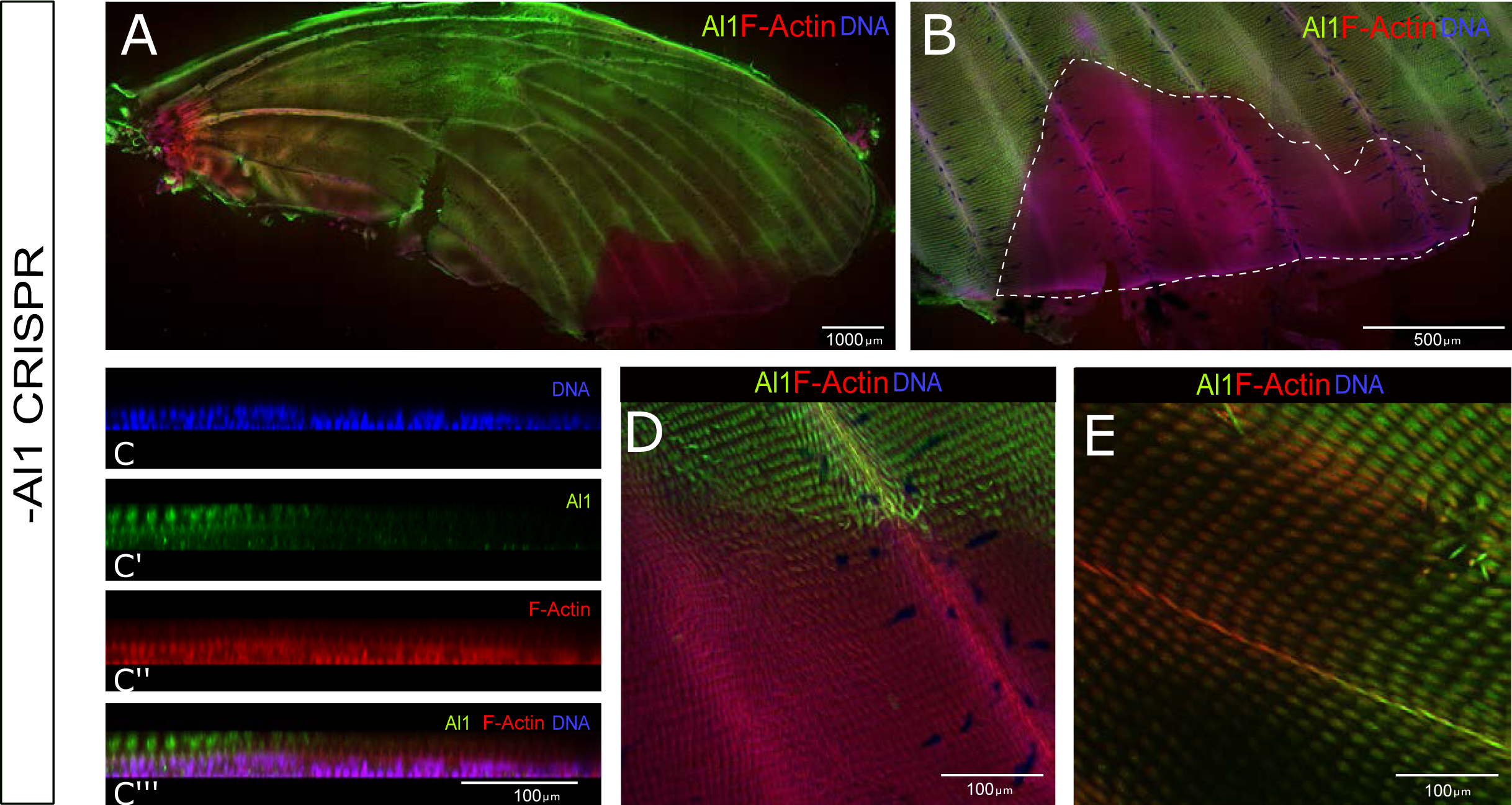
Immunodetection of Aristaless1 in *al1* CRISPR knockout pupal wings of white *Heliconius* cydno forewings (3 Days APF). (**A**) Immunodetection of Al1 in an *al1* CRISPR knockout forewing shows Al1 depleted clones. (**B**) Closer view of an extensive clone (white dashed line) within the distal edge of the wing. (**C**) Side digital reconstruction from a z stack showing the transition from high Al1 (left, outside the clone) to absent Al1 (right, within the clone). Panels show detection of DNA (C), Al1 (C’), F-actin (C’’), and a merge (C’’’) view. (**D-E**) Views of the scale precursors inside and outside of different CRISPR clones. Panel A, B, D, and E show both Al1 and F-actin.

As a complementary approach to test this hypothesis, we used electroporation mediated RNAi (Fujiwara and Nishikawa, 2016) to locally knockdown *al1* in a specific area of the wing. RNAi injections performed hours after pupation recapitulated the white to yellow color switch observed on adult wings observed previously with CRISPR/Cas9 (**Figure S8A-B**). Pupal wing discs were also analyzed by immunostaining at 3 days APF to determine if there was any effect in the protein localization of Al1 after RNAi knockdown. As expected, we found that clones with scales lacking Al1 (**Figure S8C-D**) were concentrated near the injection site. Water injection controls showed no effect on developing scale cells from the injection or electroporation process (**Figure S5E-F**). Both of these results further support our hypothesis that the white scale fate is associated with high levels of Al1 and by contrast lower levels or absent Al1 is associated with the yellow scale fate.

### Ommochrome pathway genes are differentially expressed between white and yellow wings

To infer the potential downstream consequences of differential *al1* expression, we compared expression of a number of putative pigmentation genes between white and yellow *H. cydno* wings. The difference between yellow and white wings is ultimately due to the presence or absence of the yellow pigment 3-OHK. Based on this, we focused on two enzymes involved in the production of 3-OHK, Kynurenine formamidase (Kf) and Cinnabar (Hines, et al., 2012). In addition, there is experimental evidence that 3-OHK or its precursors can be transported directly into the cell from the hemolymph (Gilbert, et al., 1988; Reed, et al., 2008). Therefore, we also analyzed the transporters White, Scarlet, Karmoisin and three members of the ATP-binding cassette (ABC) family, all of which have been implicated in 3-OHK transport or pigment movement in other *Heliconus* species (Hines, et al., 2012; **Figure 7A**). We found that the enzyme Cinnabar, as well as the transporters White, Scarlet, and Karmoisin, showed increased relative expression in yellow wings compared to white wings (**Figure 7B**). The increase in relative expression peaked at 6 days APF and exhibited the highest levels in the medial part of the wing (future yellow band). Similar differences were also observed in proximal and distal portions of the wing but to a lesser extent. Kynurenine formamidase (**Figure 7B**) and the ABC transporters (**Figure S9**) showed different trends and did not differ between white and yellow individuals. The results suggest that the white fate is achieved by reducing the expression of enzymes and transporters needed to make and move 3-OHK. This, in turn, suggests that such reduction in activity of genes needed for yellow pigmentation may be a result of Al1’s presence. We hypothesize that the reduction in Al1 expression observed earlier during pupation in yellow butterflies leads to the upregulation observed later in the enzyme Cinnabar and the transporters White, Scarlet, and Karmoisin.

**Figure 7:**
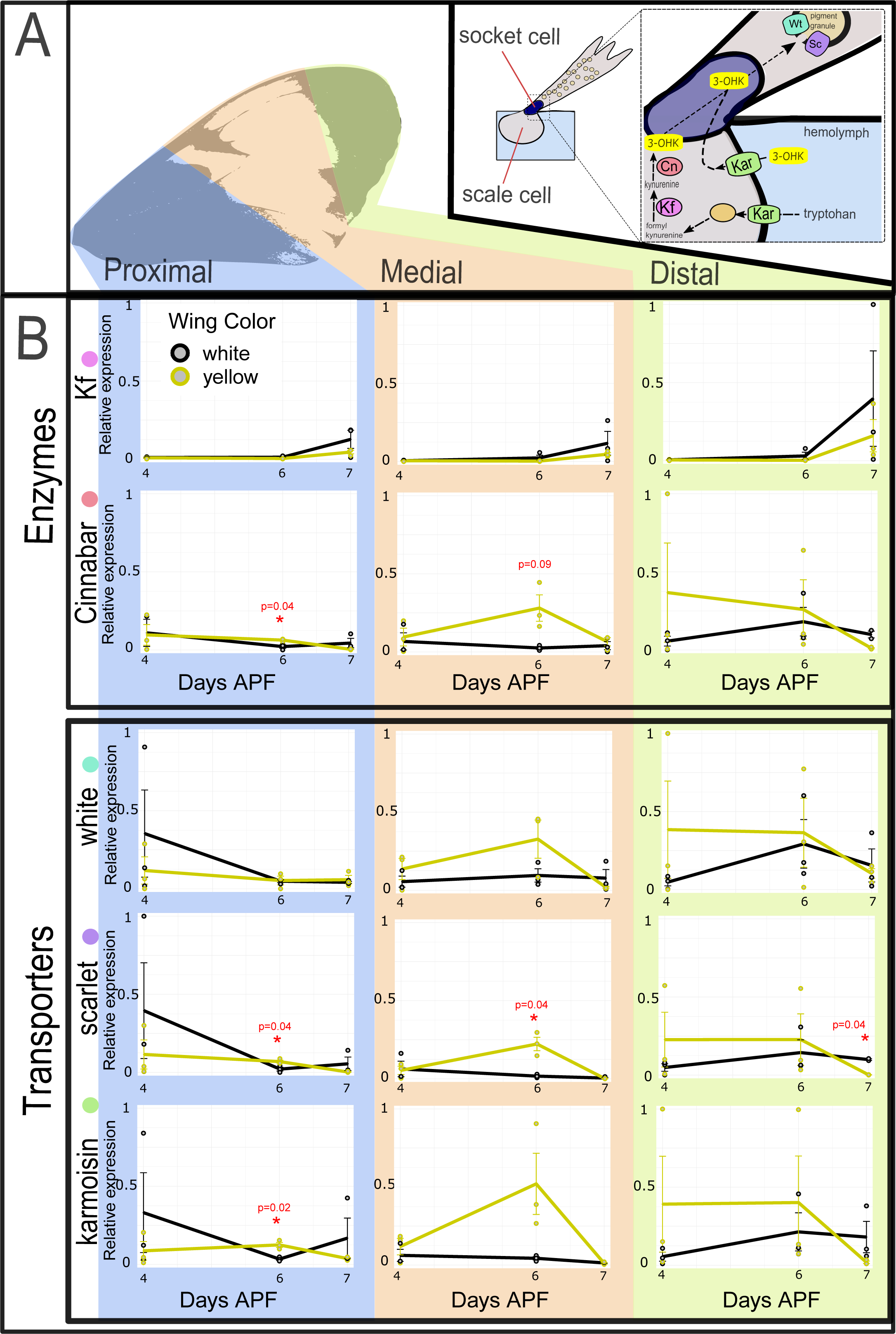
Analysis of candidate pigmentation genes that may act downstream of Aristaless1. (**A**) The top panel highlights distinct sections of the adult wing analyzed (left) and a pathway model (right) for the candidate genes of interest. The model showcases a view of the scale and socket cells and highlights the genes involved in the synthesis and transport (direct or after synthesis) of 3-hydroxykynurenine (3**-OHK**) yellow pigment. Enzymes: Kynurenine formamidase (**Kf**), Cinnabar (**Cb**); Transporters: White (**Wt**), Scarlet (**Sc**), Karmoisin (**Kar**). (**B**) Relative expression levels of each candidate gene in white and yellow pupal forewings sections (proximal, media, distal) across 3 different time points (4, 6, 7 days APF). The relative expression values are scaled to the highest value across the wing sections for each of the genes. Enzymes are shown in the middle panel and transporters on the bottom panel. Statistical significance was assessed using t-test and *p* values are shown for significant (asterisk) and nearly significant comparisons.

### Wnt signaling acts as an upstream positive regulator of Al1

Previous work on the role of Al1 in the development of moth antennae has shown that its expression is upregulated by Wnt signaling (Ando, et al., 2018). Therefore, we sought to test the potential role of Wnt signaling in the regulation of Al1 on developing *Heliconius* wings. Given that the presence of Al1 results in white scale development, we hypothesized that inhibiting Wnt-mediated transcription should lead to reduced or absent Al1 and a white to yellow switch. (**Figure 8A**). In addition, we validated our manipulations on Wnt signaling in yellow butterflies by using an inhibitor against GSK3 which should activate Wnt signaling. Because Al1 is naturally downregulated in yellow butterflies, we hypothesized that activation of Wnt signaling should enhance Al1 expression and promote a yellow to white color pattern switch (**Figure 8A**). Finally, as proof of concept that our pharmacological agents were affecting Wnt signaling, we also assayed the effects of inhibiting and activating Wnt signalizing on the development of melanic scales, which is known to be under the control of WntA activity (Martin, et al., 2014). It has been shown that scales lacking WntA activity become paler or completely revert to a different color fate from the wing (Mazo-Vargas, et al., 2017). Furthermore, previous work has shown that increasing Wnt responsive activity in non-melanic parts of the wing by using the pharmacological agent Heparin switches non-melanic scales into melanic ones (Martin, et al., 2012). Therefore, we hypothesized that reduced Wnt activity in melanic portions of the wing should result in paler or non-melanic scales while activating Wnt in non-melanic parts of the wing should promote melanization (**Figure 8A**).

**Figure 8:**
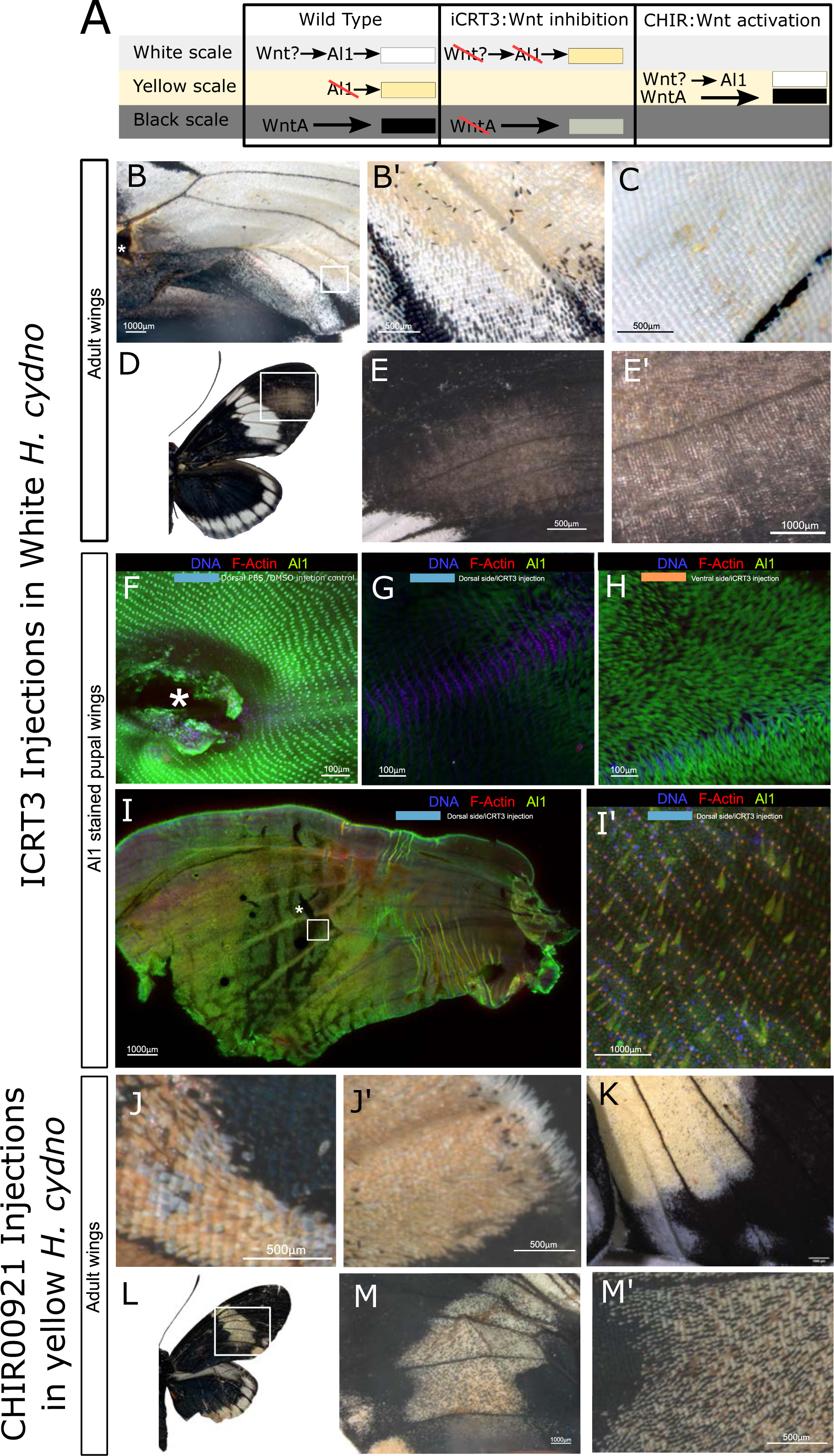
Aristaless1 is regulated by Wnt signaling. (**A**) Scheme for the phenotypic color outcome for both wild type white and yellow butterflies. The hypothesized scenarios caused after exposure to the iCRT3 and CHIR00921 inhibitors affecting Wnt signaling is presented for both the white and yellow background. In it, we expect the white to yellow color switch following the reduction in Al1 levels caused by diminished Wnt signaling and the yellow to white switch following increase Al1 as a cause from enhancing Wnt signaling. Outcomes with respect to melanic scales are also shown as a read out from WntA activity. (**B**) Adult white wing injected at 3 days APF with iCRT3 and observed after eclosion. White square is shown as a detail view in C’. (**C**) Adult wing on the opposite side to the injection. (**D-E**) Adult right wing showing the effects of iCRT3 on melanic scales. Detailed view of the affected side (**E**) and scales are shown (**E’**). (**F-I**) Developmental validation of the iCRT3 effects on Al1 protein levels. The injection control (**F**) with 1Xpbs/DMSO is show as well as the dorsal (**G**) and ventral (**H**) sides of a pupal wing exposed to the drug around 3 days APF and dissected 24 hours after exposure to the agent. A different full wing with the same treatment is shown (**I**) with a wider area of effect. Severe scale size defects are visible in an amplified view from the white square (**I’**). (**J**) Different parts of an Adult yellow wing injected at 3 days APF with CHIR00921 and overserved after eclosion. (**K**) Ventral side of a different individual treated in the same way. (**L-M**) Adult right wing showing the effects of CHIR00921 with respect to the formartion of melanic scales. Details are shown (**M**). Asterisk showcases injection sites. In F-G the injection site is on the left outside the field of view.

Our data showed that exposing the pupal wing to the Wnt signaling inhibitor iCRT3 did produce a white to yellow switch as predicted (**Figure 8B-C**). In parallel, when the Wnt inhibitor was used on melanic parts we observed the change from black to a paler color as expected from a WntA knockdown (**Figure 8D-E**). Furthermore, wings exposed to the inhibitor also showed depleted levels of Al1 when comparing the dorsal (in closer contact to iCRT3) and ventral sections on the wing (**Figure 8E-G**). DMSO/PBS controls showed normal Al1 levels, highlighting that the procedure itself did not cause the observed effect (**Figure 8F**). Furthermore, the untreated wing of the same butterfly showed normal levels of Al1 as well. Yellow wings that were treated with the GSK3 inhibitor CHIR99021, which promotes Wnt signaling, developed white scales as hypothesized (**Figure 8J-K**). Finally, we also observe several melanic scales within yellow band region as expected by a Wnt gain of function (**Figure 8L-M**).

Following exposure to iCRT3, some wings exhibited zones with peculiar scale phenotypes (**Figure 8H**). Examination of these zones showed that some of the scales were normal size and had normal Al1 levels but others were smaller and exhibited lower Al1 levels (**Figure 8H**’). To our knowledge, there have not been any reports of scales showing differential growth rates within the same scale fate. This may be a secondary effect from other gene targets affected by inhibited Wnt signaling and then the lower Al1 levels are just a result of a smaller scale. An alternative explanation could be that Al1 also influences processes related to scale growth and elongation (as shown in other systems; Campbell and Tomlinson, 1988; Schneitz, et al., 1993; Ando, et al., 2018) and by partially depleting its levels with iCRT3 we are altering those functions.

## Discussion

Our results suggest a model for how the decision between white and yellow scale fate is achieved under the control of *al1* during wing development in *Heliconius* butterflies (**Figure 9**). Overall, our data show that Al1 accumulates within the cytoplasm of future white and melanic scales but is depleted from future yellow scales during the early stages of pupation (2 days APF). These results suggest that the presence of Al1 within the cytoplasm is relevant for the specification and/or pigmentation of both white and black scales but not yellow scales. Evidence in favor of this model includes *al1* CRISPR/Cas9 knockout clones that span both white and black portions of the wing.

**Figure 9:**
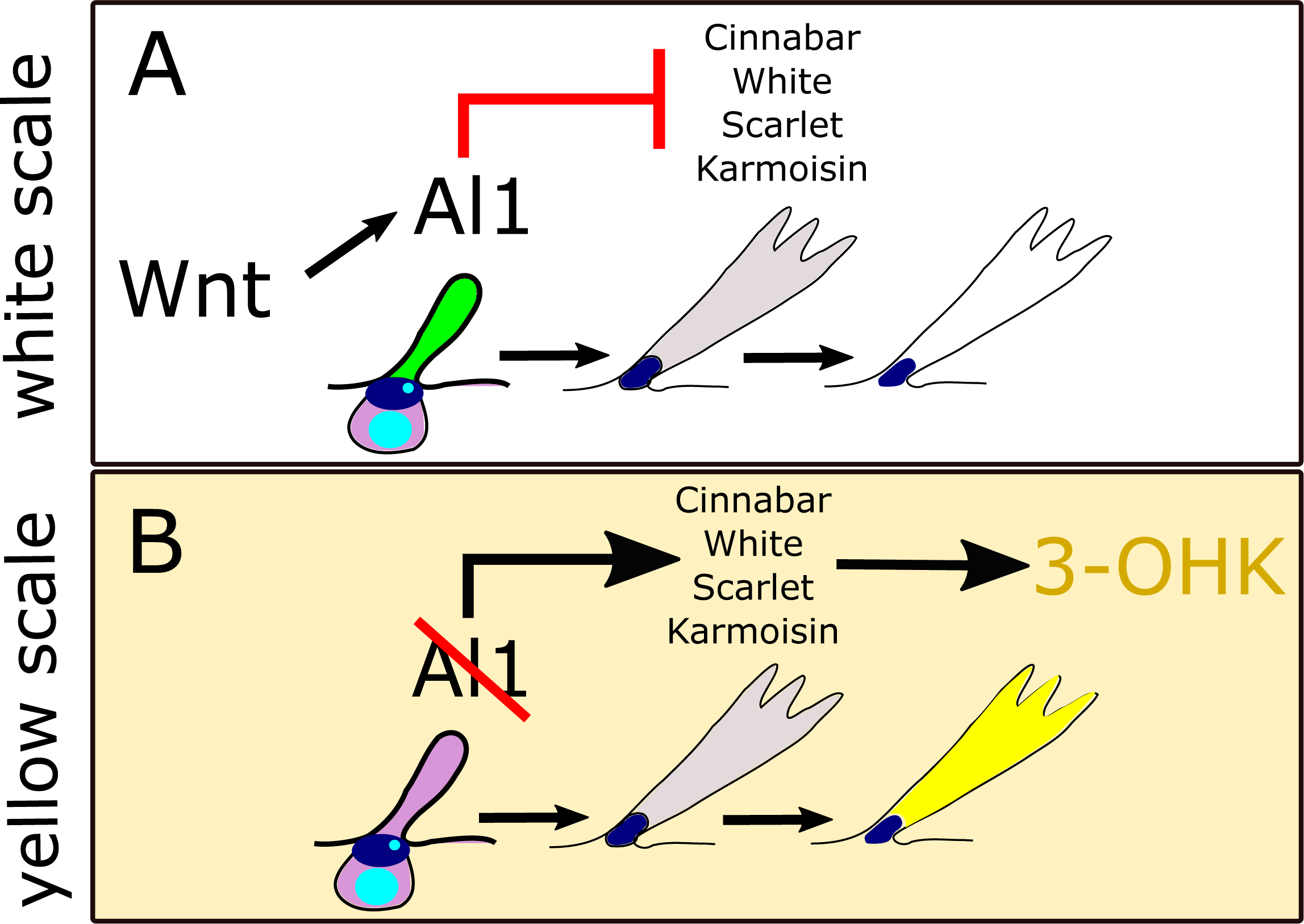
Graphical model for the role of Al1 in the specification *H. cydno* wing color. (**A**) White scale fate in which Al1 presence in developing scale cells lead to the inhibition of genes needed for yellow pigment uptake and production. (**B**) Yellow scale fate in which, reduced or absent Al1 results in the activation of genes involved with the production and movement of the yellow pigment 3-OHK.

Scales within these clones show a switch to yellow and brown respectively. However, *al1* knockouts have no observed effects within the yellow band. Knockouts by CRISPR/Cas9 and knockdowns by RNAi result in depleted levels of *al1* in developing scales during early pupation as well as an associated switch from white to yellow scales. Our model is further informed by the preliminary observation that Al1 seems to promote the white color fate by negatively regulating genes important for the synthesis and transport of 3-OHK. In addition, we also validated the role of Wnt as an important upstream signal for Al1 activation providing a more complete developmental context. These functional data highlight how Al1 specifies the development of black and white scales and inhibits yellow pigmentation.

Our results for *aristaless1*’s role in the control of white and yellow wing coloration provide a different patterning scheme for the specification of wing color patterns. Previous work with other *Heliconius* color patterning genes has shown how the expression of these genes during earlier developmental stages (larval or pupal) resembles the future adult color pattern (Reed, et al., 2011; Martin, et al., 2012; Martin, et al., 2014; Nadeau, et al., 2016). This spatial prefiguring is very clear with all three of the previously described *Heliconius* color patterning genes: *optix* (Martin, et al., 2014), *wntA* (Martin, et al., 2012) and, *cortex* (Nadeau, et al., 2016). Furthermore, CRISPR/Cas9 knockouts of both *optix* (Zhang, et al., 2017) and *wntA* (Mazo-Vargas, et al., 2017) result in the lack of their respective color patterns. All of these genes, acting as activators, organize and promote their respective color patterns. On the other hand, we observe that Al1 is present in the entire wing and represses the yellow scale fate. It is the absence of that repression which ultimately results in the color switch and pattern establishment we observe in the adult. While repression is a well-described developmental phenomenon, the color pattern variation achieved via repression of *al1* makes this a unique mechanism relative to other *Heliconius* color patterning genes.

Considering *al1* along with *wntA*, *optix*, and *cortex* it becomes clear that even though all of these genes control wing color patterning, they do so by very different mechanisms. For example, WntA is a signaling ligand that has its effect early within the larval imaginal discs (Martin, et al., 2012). As a signaling molecule WntA is restricted in its ability to diffuse to other nearby cells (Martin, et al., 2012) and therefore it may function primarily during larval development as opposed to pupal development where scale cells are more discrete and distantly distributed. Optix is a transcription factor that is directly localized to the nucleus of red scale precursors during mid-pupation (Martin, et al., 2014), possibly activating downstream targets needed to eventually produce red scales. Cortex is another unique scenario; as a cell cycle regulator in other systems, it is currently unknown how such a protein controls the melanic color patterns it resembles during its pupal expression (Nadeau, et al., 2016). Finally, Al1 is a homeodomain protein involved in appendage extension (Campbell and Tomlinson, 1988; Schneitz, et al., 1993; Beermann and Schroder, 2004, Ando, et al., 2018) which we have found to control multiple aspects of wing pigmentation. Al1 does this by localizing within scale cell precursors during early pupation yet it is specifically depleted from future yellow scales. This information highlights that very different types of genes can be major regulators of color patterning by employing various mechanisms associated with their identity. This developmental description of *al1* serves as the foundation for trying to answer the question of how differences in the levels of *al1* result in the white and yellow color switch. Here we have provided evidence In favor of a model whereby *al1* is, by some direct or indirect mechanism, acting as a repressor of genes involved in yellow pigmentation.

In terms of a direct mechanism, the most straightforward scenario involves Al1 repressing genes involved in yellow pigmentation (*cinnabar*, *white*, *scarlet,* and *karmoisin*) in the nucleus, as expected of a transcription factor. However, we are particularly intrigued by the observation that Al1 was never found localized in the nucleus during the analyzed time points. It is important to acknowledge that there could still be a specific time point in which Al1 translocation happens leading to the transcriptional control of downstream proteins needed for proper yellow pigmentation. In addition, there is a possibility that a post-translational modification—for example a cleavage event like the ones observed in BMPs proteins (Künnapuu, et al. 2009) or in another Paired-like homeodomain protein ESXR1 (N-terminus translocate to the nucleus and C terminus stays cytoplasmic, Ozawa, et al., 2004)—occurs with Al1 which affects our ability to observe nuclear co-localization. However, regardless of the possibility of our inability to observe a possible nuclear localization of Al1, there is still experimental evidence showcasing that some transcription factors can regulate other downstream processes and showcase dynamic states between cytoplasmic and nuclear localizations. For example, the protein Extradentricle (Exd) which is exported to the cytoplasm when Homothorax is not present (Abu-Shaar, et al., 1999) can exhibit different patterns of cytoplasmic or nuclear localization depending on what part of the leg imaginal disc is being patterned (Abu-Shaar, et al., 1999). Furthermore, in such a system an increase in the accumulation from cytoplasmic Exd can lead to an overcoming of the signals keeping the protein cytoplasmic, allowing a portion of them to go into the nucleus even when Homothorax is not present (Abu-Shaar, et al., 1999). This is an interesting case considering that both Exd and Al1 are homeodomain proteins and similar accumulation is visible in our data. Therefore, it is possible that Al1 could act as a direct regulator (by an un-observed nuclear translocation or a cleavage event) of the differentially expressed genes needed for yellow pigmentation.

An alternative possibility is that Al1 regulates wing pigmentation indirectly via a process known as the Heterochrony hypothesis (Koch, et al. 2000). This is an interesting possibility based on what we know about the role of Aristaless in appendage extension (Campbell and Tomlinson, 1988; Schneitz, et al., 1993; Ando, et al., 2018) and based on our data showing dIfficiency of appendage extension following Al1 knockouts. Although, Aristaless is described as a homeodomain transcription factor, most of the literature describing its expression and subcellular localization is related to its role during the extension of body appendages at both the single-cell and multicellular level (Campbell and Tomlinson, 1988; Schneitz, et al., 1993; Ando, et al., 2018). In *Drosophila*, Aristaless is well characterized for its role in the extension and proper patterning of the arista (a highly modifiable bristle that extends out of the antennae). Previous work has shown that if Aristaless is not present, pronounced size reductions and malformations of the arista occur (Schneitz, et al., 1993). Similar elongation defects to the ones we observed in our embryos are seen when Al1 expression is reduced in the multicellular antennae of moths. In this system, Al1 is needed for the proper patterning and the directional elongation of the cells that form part of the antennae. Furthermore, outside of insects the Aristaless-like Homeobox (ALX) protein is a key regulator of rodent pigmentation (Mallarino, et al., 2016). Such regulation in principle is controlled by adjusting the rate of maturation of melanocytes, which are the pigmented cells that ultimately carry out the pigment synthesis of the hairs on the rodent body (Mallarino, et al., 2016). These observations support the idea that Al1 could be controlling pigmentation outcomes by altering rate of scale development. Another, piece of evidence that further promotes Al1 as a candidate capable of regulating the cell cycle and affecting scale maturation time, is again the Paired-like homeodomain protein ESXR1. The C-terminal region of ESXRI stays in the cytoplasm after proteolytic cleavage and inhibits cyclin degradation which regulates the cell cycle and even produces cellular arrest (Ozawa, et al., 2004). This effect on the cell cycle produced by a cleaved component of a paired-like homeodomain protein makes it an appealing mechanism for the heterochronic shift we are hypothesizing. These examples raise the possibility that Al1 may be altering the developmental rate of scales which, in turn, influences color by indirectly altering expression windows of transporters and enzymes necessary for pigmentation. Yellow pigmentation in *Heliconius* happens just a few hours before eclosion, and therefore small alterations to the developmental timing of scales could result in the presence or absence of 3-OHK.

Future work will determine whether Al1 directly affects downstream target genes by regulating their transcription or indirectly as a secondary effect from altering scale maturation time. Our work serves as the first developmental description of Al1 and helps us understand butterfly color patterning as a stepwise process involving multiple layers of gene regulation terminating in pigmentation. Our work also highlights the diversity of genes and developmental mechanisms responsible for butterfly wing patterning.

## Methods

### Butterflies rearing

Butterflies were reared in greenhouses at the University of Chicago with a 16h:8h light:dark cycle at ∼27°C and 60% – 80% humidity. Adults were fed Bird’s Choice artificial butterfly nectar. Larvae were raised on *Passiflora biflora* and *Passiflora oerstedii*.

### CRISPR/Cas9 injections

CRISPR/Cas9 experiments were performed following Westerman et al. (2018). We used HC_gRNA_02_Al1 (GTTCTAGGAGAATCGTCCTTTGG) and HC_gRNA_03_Al1 (GGAGGAGGTCTCTCGGAGGCTGG) gRNAs to generate deletions in *Al1* in *Heliconius cydno galanthus* and *Heliconius cydno alithea* (**Figure 2**). The concentration of Cas9 (PNA Bio) and sgRNAs varied between 125 ng/µl−250 ng/µl and 83 ng/µl−125 ng/μl, respectively. Mixes were heated to 37°C for 10 min immediately prior to injection and kept at room temperature while injecting. To collect eggs for injections, we offered adults fresh *Passiflora oerstedii* and allowed ∼2 hours for oviposition. Eggs were washed for 2 min in 7.5% benzalkonium chloride (Sigma Aldrich), rinsed thoroughly with water, and then arrayed on double-sided tape on a glass slide for injection. The eggs were injected using a 0.5-mm borosilicate needle (Sutter Instruments, Novato, CA, USA) and then kept in a humid petri dish until hatching, then transferred to a fresh host plant and allowed to develop. Adults were frozen and pinned before imaging. Following injection, 69 white and 4 yellow individuals reached adulthood. From them, 40 white, and 3 yellow individuals had a phenotype.

### Imaging of wild type and CRISPR adult wings

Butterflies were pinned to flatten the wings and dry the tissue allowing for better imaging and then photographed. Details of wild type and adult wings were imaged using a Zeiss stereomicroscope Discovery.V20 with AxioCam adapter. Z-stacks and maximum intensity projections were produced using the Axiovision software. All Images had their intensity and scale bars edited with ImageJ Software.

### Butterfly wing dissections

Butterflies were dissected at both larval and pupal stages following Martin et al. (2014). The protocol and adaptations to it were carried out as follows. Larvae and pupae were anesthetized in ice for 20 mins before dissection. To obtain the larval wing discs the larvae were pinned on the first and last segment. A small cut was performed using micro-dissection scissors on the second (forewing) and third segment (hindwing) to remove the imaginal discs. The discs were then pipetted out to a 16 well tissue culture plate with 1 ml per well of a 4% Paraformaldehyde solution for fixing. Larval imaginal discs were then fixed between 20 to 30 mins. To obtain the pupal wing discs the pupae were pinned on the head and most posterior section of the body. The denticle belt was then removed using dissection forceps to allow for easier access to the wing. Then micro-dissection scissors were used to carefully cut around the wing margin using the pupal cuticle as a guide. The piece of cuticle together with the forewing imaginal disc was removed and placed directly in a 16 well tissue culture plate with 1 ml per well of a 4% Paraformaldehyde solution for fixing. Pupal wings were fixed for 30 to 45 mins and then cleaned of any peripodial membrane by using fine forceps. After fixation, the tissue was then washed with PBST (PBS + 0.5% Triton-X100 for antibody staining or with PBS, + 0.01% Tween20 for *in situ* hybridization) five times to then be stored at 4°C until stained (not more than 30 days).

### Embryos fixation and dissection

Eggs were collected from plants between 24 to 36 hours after deposition. We adapted the fixation scheme from Brakefield et al. (2009). Eggs were first transferred to 1.5 ml tubes and washed on PBS to remove any dirt. Eggs were then permeabilized and had their chorion removed with 5% Bleach (PBS) for 6 minutes. Eggs were then washed 5 times for 5 minutes in PBS to remove the excess bleach. We added 1 ml per tube of a 4% Paraformaldehyde solution (PBS) for fixing for 30 to 60 minutes. Eggs were then washed in PBST (PBS + 0.5% Triton-X100) 2 times for 5 minutes and then taken into a methanol series (25%, 50%, 75% methanol solutions in PBS at 4◦C). Eggs were then transferred to 100% methanol and stored at -20◦C for 5 days. Eggs were then transferred using plastic pipettes to a glass dissection plate with pre-chilled 100% methanol for dissection with fine forceps and dissection needles. Dissected embryos were then pipetted carefully into a 16 well tissue culture plate with 1 ml per well of chill methanol. These embryos were taken back through a 1 ml per well methanol series (75%, 50%, 25% methanol solutions in PBS at 4◦C) for rehydration. Then embryos were washed twice with 1 ml of PBST per well and stored in PBST at 4◦C for antibody staining.

### *al1* in situ hybridization of larval wings

We designed and synthesized *al1-*specific probes using the *H. cydno al1* transcript model (selected region shows 100% identity with *aristaless1* and 60% identity with *aristaless*2 transcript model). A 250 base-pair region from *al1* was amplified using primers (forward GTTCCCTCGCAGCCATTCTT; reverse TACGGCACTTCACCAGTTCT) by PCR, cloned into a TOPO vector (Invitrogen), and transformed into competent *E. coli* DH5a cells. We grew 3 replicates of 2 positive colonies and extracted DNA using a miniprep DNA extraction kit. We confirmed insert sequences via Sanger sequencing, linearized plasmids using Not1 and Sac1 restriction enzymes (New England Biolabs), and synthesized probes using a reverse transcription kit (Qiagen) with added DIG labeled nucleotides. The synthesized probes were purified using Qiagen RNAeasy columns.

*In situs* were performed following Ramos and Monteiro (2007). The entire process was carried out in 16 well tissue culture plates. Tissues stored in PBST (PBS, Tween20) were subjected to a mild digestion for 5 minutes in Proteinase K (0.025mg/ml). Digestion was stopped using a stop buffer (2mg/ml glycine in PBS 0.01% tween20). Tissue was washed 5 times for 5 min with PBST, then incubated in a pre-hybridization buffer (50%formamide, 5XSSC, 0.1% Tween20, and 1mg/ml Salmon Sperm DNA) for 1 hour at 55°C. 1 ml of Hybridization buffer (50%formamide, 0.01g/ml glycine, 5XSSC, 0.1% Tween20, and 1mg/ml Salmon Sperm DNA) with approximately 50 ng of the used probe against *al1* was added to each well and left to incubate at 55°C for at least 48 hours. The tissue was then washed 5 times for 5 min in pre-hybridization buffer and then left washing in pre-hybridization buffer for 24 hours at 55°C. Wings were then blocked in 1% bovine serum albumin (BSA) in pre-hybridization buffer for 1 hr at 4°C. Anti-DIG antibody was added (1:2000) to each of the wells and incubated overnight at 4°C. The tissue was then washed with PBST extensively (10 times or more for 5 minutes) before development with BM-purple (1ml peer well, Roche Diagnostics). Time of development was approximately 24 hours at 4°C. Stained tissue was imaged using Zeiss stereomicroscope Discovery.V20 with AxioCam adapter. Scale bars were added using ImageJ software. We analyzed wing imaginal discs of white butterflies at both fourth and fifth instar stages (3 individuals, wings split between sense and antisense probes).

### Al1 antibody staining of embryos, larval, and pupal wings

We raised polyclonal antibodies against two Al1 peptides using GenScript (New Jersey, USA). Peptide antigens (Al1-1: QSPASERPPPGSADC, Al1-2: DDSPRTTPELSHA) are located in the N-terminal 40 amino acids of Al1 and share 25% and 30% identity with Al2. Polyclonal antibodies were affinity purified after harvesting and tested for specificity by performing Dot blot tests as described in **Figure S2**. Raw data is available in **Source Data 1-2.**

We performed antibody staining in larval and pupal wings following Martin et al. (2014). We also applied this staining protocol to embryos. Tissue stored in PBST (PBS, Tritonx) was blocked in 1% BSA in PBST for two hours, then incubated overnight in 1 mL blocking buffer and Al1 specific antibody (1:1000 for pupal wings and embryos, 1:300 for larval wings). Tissue was washed twice quickly, then 5 times for 5 mins in ∼0.5 mL PBST, then incubated in 1 mL the secondary staining solution (goat anti-rabbit-AlexaFluor 488 [Thermofisher] at 1:1000, Hoechst 33342 at 1:1000 [Thermofisher] and Phalloidin-AlexaFluor555 at 1:200 [Thermofisher] in blocking buffer). The tissue was washed extensively and then mounted on glass slides using VectaShield (Vector Labs) on glass slides. Images were collected using a Zeiss LSM 710 Confocal Microscope and processed using Zen 2012 (Zeiss) and ImageJ. For wild type antibody stainings we used pupal wings between 2-4 days APF of both white and yellow butterflies (20 individuals for white and 6 individuals for yellow). For white CRISPR knockout butterflies we used wings 2 days APF, (3 individuals, 2 of which showed a phenotype), 3 days APF (4 individuals, 3 of which showed a phenotypes), and 4 days APF (3 individuals, 2 of which showed a phenotype). For embryos we used 5 wild type and 4 CRISPR embryos (3 of which had a phenotype).

### Electroporation of pupal wings for RNA interference

Electroporation-mediated RNA interference experiments were performed following Ando and Fujiwara (2013) and Fujiwara and Nishikawa (2016). We designed and synthesized Dicer substrate short interfering RNAs (DsiRNAs) targeting the first exon of *Al1* using Integrated DNA Technologies (USA). Al1.DsiRNA-1 targets 5’-ATGAATTTACTCCAAAAAGAAAG.

Fresh pupae, within the first hour of pupation, were used to perform the injections. For each experiment, the pupa was placed on a petri dish under a stereoscope and had its forewing displaced over a 1% agar (1xPBS) pad. One microliter of 250 µM DsiRNA in water was injected into one of the pupal wings using borosilicate glass needles (with filament; GDC-1 from Narishige, USA) pulled on a Narishige PC-10 with 1 step at setting 67. A 1xPBS bubble was placed on top of the injection site to perform electroporation using 5 x 280 ms pulses of 10 V over 5 sec. The wing was then placed back in its original position and the insect was allowed to recover for 24 hours before being hung again vertically. Some electroporated pupae were allowed to develop to ad ulthood and others were dissected 3 days APF for staining following the methods described above. Approximately 45 pupae were treated. We used wings at 3 days APF from 5 individuals for Al1 stainings (3 of which showed a phenotype). From the remaining 40 pupae, 14 survived to at least pre-eclosion stages (5 showed an adult phenotype).

### qPCR gene expression analysis of downstream target genes

We collected pupal forewings 4, 6, and 7 days APF of both white and yellow *Heliconius cydno* butterflies (three biological replicates of each color at each time point). The collected wings were cut into 3 sections (proximal, medial, and distal) using the venation pattern as a guide for consistent cuts (**Figure 8A**). Following dissection, the tissue was stored in RNA later (Ambion, USA) at -80°C until RNA extraction. The same sections from the two wings in each individual were grouped. Samples were thawed on ice, then washed twice with ice cold PBS before total RNA extraction using TRIzol (Invitrogen, USA) and the manufacturer’s protocol. Extracted RNA was re-suspended in 50 µL of RNAse free water. Purified RNA (2 µg) was used to perform cDNA synthesis using the ABI High Capacity cDNA Reverse Transcription Kit (Thermo Fisher 4368814) following the manufacturer’s instructions. cDNA pools were diluted 10X in TE and stored at 4°C until qPCR.

All qPCRs were performed in 10 uL reactions with the BioRad iTaq Universal SYBR Green Supermix (Bio-Rad, USA) on a Bio-Rad CFX96 thermal cycler. We tested primer efficiencies using a 2-fold dilution series of one cDNA pool and only used those with efficiencies between 90% and 106% when possible (**Table S1**). We used *ef1a* as the ubiquitously expressed control gene to standardize our values during the qPCR assays. A single experimental gene and the control gene were tested for all samples in a single plate, and all reactions were technically replicated twice. Relative expression levels were calculated using the ΔC_T_ method. We then scaled ΔC_T_ values for a particular gene to 1 by dividing sample ΔC_T_ values by the highest ΔC_T_ value for that gene. Calculations were performed in R (version 3.5.2). Raw data is available in **Source Data 3**.

### ICRT3 and CHIR99021 test on Wnt signaling

The inhibitor of β-catenin responsive transcription (iCRT3, MedChemExpress Cat. No.: HY-103705, stock concentration; 10 µg/µL in DMSO) and the inhibitor of the repressor GSK3 α/β, (CHIR99021, Sigma-Aldrich Cat. No.: 252917-06-9 stock concentration; 5 µg/µL in DMSO) were used on pupal wings 2 to 4 days APF. The pupae were cold anesthetized for 5 minutes before making a small opening on the cutical covering of the pupal wing. Then the piece of cuticle covering the opening was lifted in order to add 3µL of the inhibitor solution (400ng/µL iCRT3/ CHIR99021 in PBS1x or in 1xPBS/DMSO). For controls, pupae with just the opening as well as pupae with 3 µL of 1XPBS/DMSO were used. After the addition of the solutions, the piece of the cuticle was placed back and the pupa was left resting without hanging for 24 hours to allow for healing and recovery. If the wing was going to be imaged the dissection and staining was carried out as described above. In the case where the butterfly was going to be allowed to develop to adulthood it was hung again between 24 to 48 hours after exposure and taken back to the greenhouse. Approximately 60 pupae of white *H. cydno* were treated with ICRT3. We used wings between 2 to 4 days APF from 10 individuals for Al1 antibody stainings (6 of which showed a phenotype [2 of them had scale size phenotypes]). Of the remaining 50 treated butterflies, 15 survived to at least pre-eclosion stages (7 showed an adult phenotype). Three yellow H. cydno pupae were treated with CHIR99021, of which all 3 showed one or both phenotypes of yellow scales switching to white or black.

## Acknowledgments

We thank Michael Hennessy and Carlos Sahagun for butterfly care and Steven Lane for assistance with dissections and staining. We also thank Urs Schmidt-Ott, Victoria Prince, Stephanie Palmer, and reviewers for discussion and/or comments on the manuscript.

## Supplemental Figures

**Supplemental Figure 1:**
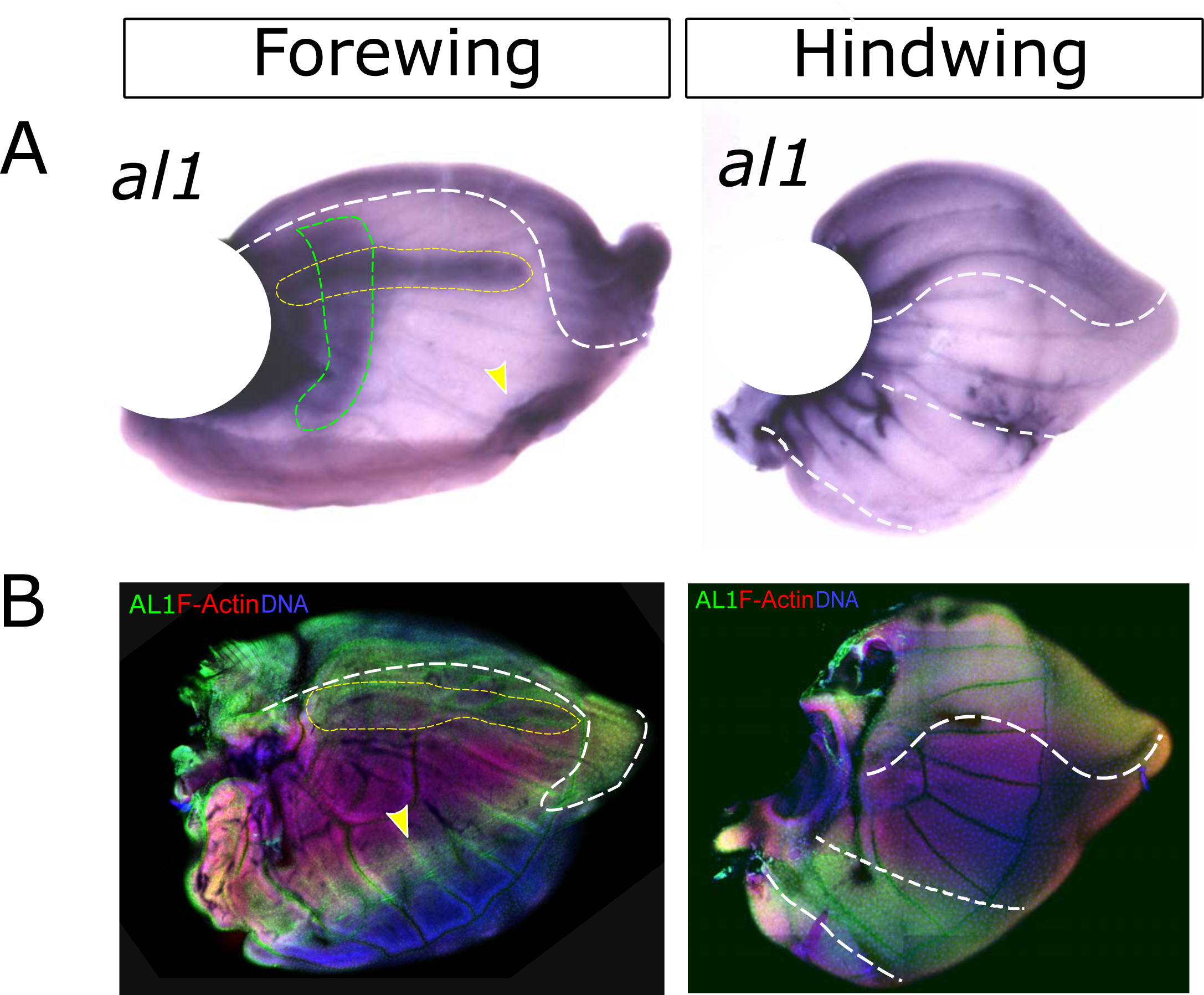
Detection of aristaless1 by mRNA *in situ* hybridization and Al1 specific antibodies in white *H. cydno*. (A) mRNA *in situ* hybridization of 5th instar larval forewing and hindwing. (B) Al1 antibody staining of 5th instar larval forewing and hindwing. Dotted lines are used to highlight previously described domains of expression from Martin and Reed (2010). White dotted lines showcase the anterior curved domain (both forewings and hindwings) and posterior narrow band (hindwings). The yellow dotted lines highlight the horizontal expression domain along the anterior veins of forewings. The green dotted line highlights a vertical expression domain observed only in our *in situ* hybridization forewing. This domain has previously been reported as an Al2 expression pattern, suggesting some cross-reaction from the used probe. The yellow arrowhead highlights a posterior expression domain not previously described before and observed in both *in situ* and antibody-stained forewings.

**Supplemental Figure 2:**
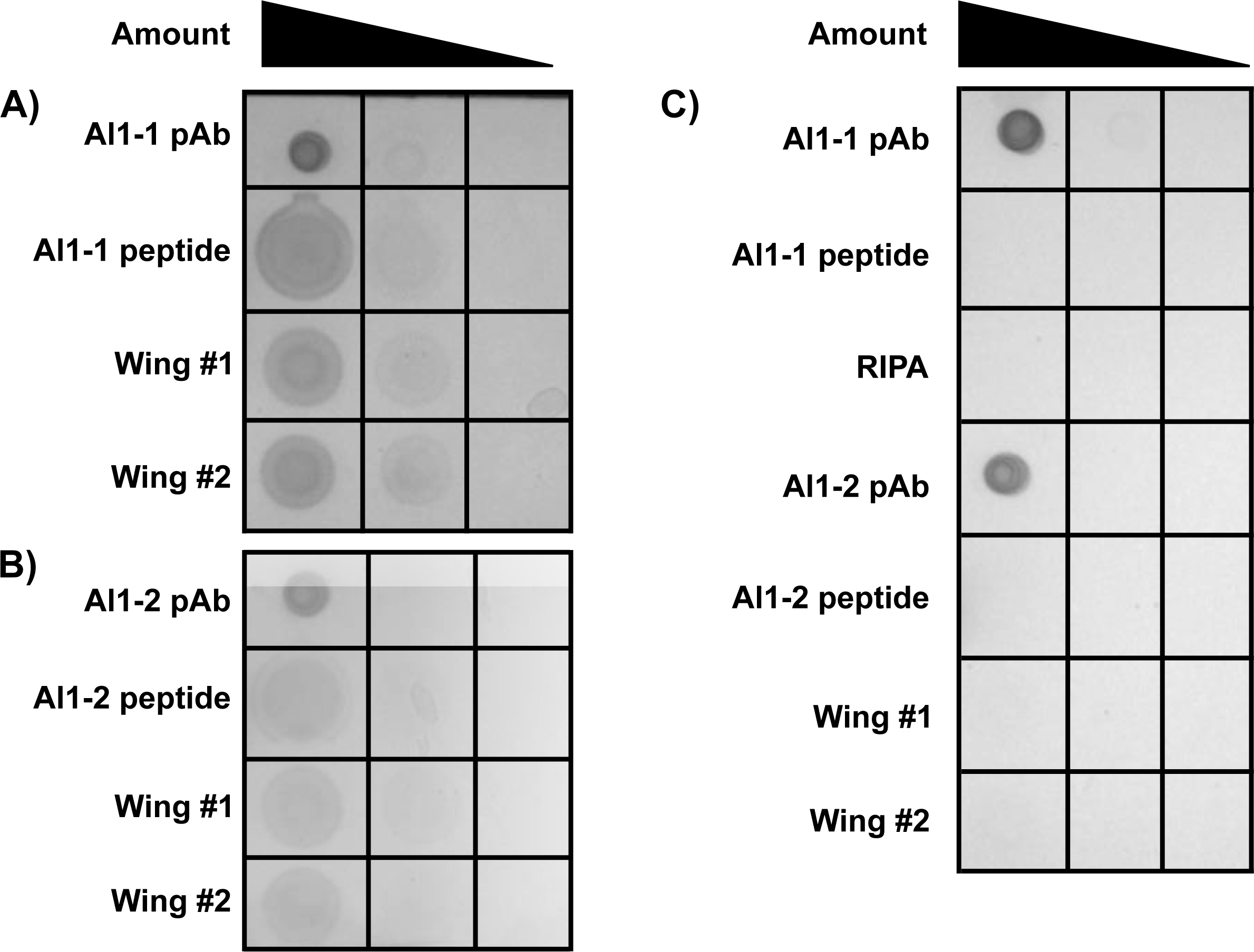
Dot blot test to determine the specificity of the Al1 antibodies. We spotted 2 uL each of three amounts of each antibody, peptide antigen, or protein prep (Wing #1, Wing #2), then probed blots using 5 ug/mL Al1-1 (**A**), 5 ug/mL Al1-2 (**B**), or no primary antibody (**C**). All blots were then probed with goat anti-rabbit secondary antibody conjugated to alkaline phosphatase (Invitrogen 65-6122). All three blots were developed for 15 min in the same container using Roche BM Purple AP substrate (11442074001) before imaging on a BioRad GelDoc XR+. Dot amounts: antibodies and peptides = 200 ng, 20 ng, 2 ng; protein preps: 1X, 0.2X, 0.05X.

**Supplemental Figure 3:**
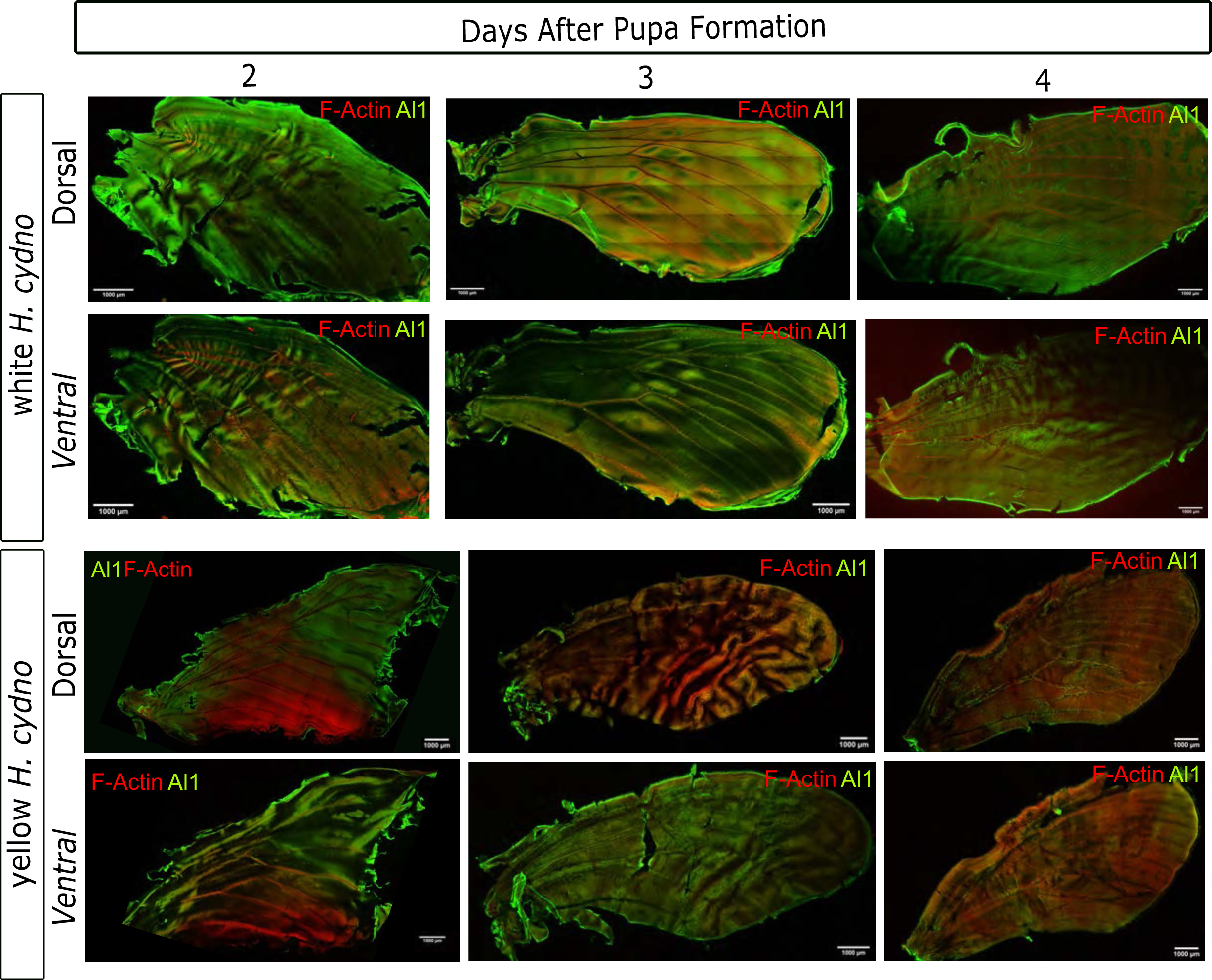
Temporal and spatial differences in Al1 protein localization between white and yellow *Heliconius cydno* wings. (A) Immunodetection of Al1 in white *H. cydno* forewings at different stages of early pupation (2 to 4 days APF) for both the ventral and dorsal side of the wing. **(B)** Immunodetection of Al1 in yellow *H. cydno* forewings at comparable stages to the white wings in panel A. Both ventral and dorsal parts of the wing are shown as well. Both panels show detection of Al1 and actin.

**Supplemental Figure 4:**
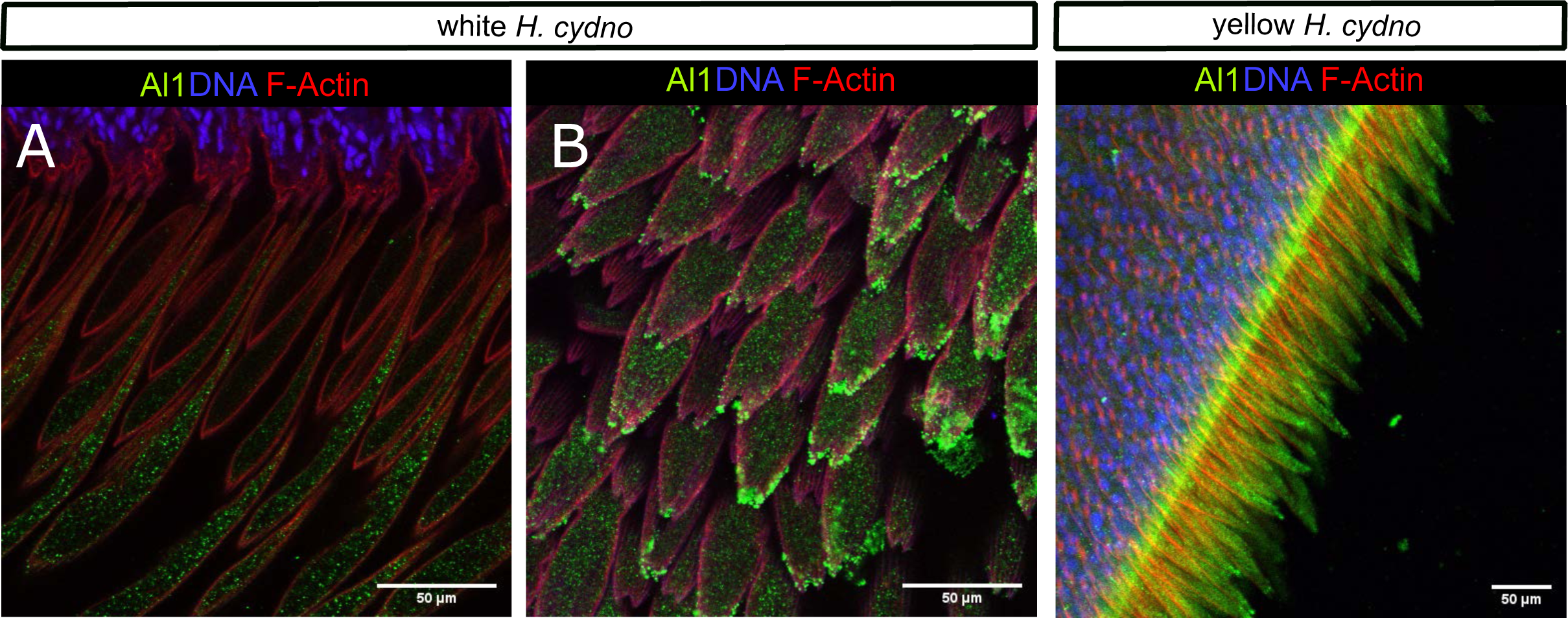
Immunodetection of Aristaless1 in melanic scales for both white and yellow *Heliconius cydno* pupal forewings (late 4 Days APF). (**A**) Imaging of longer border scales to appreciate details on the protein subcellular localization. View of bi-forked (**B**) and tri-forked (**C**) scales with accumulating Al1 in the scale cell body of a yellow *H. cydno* highlighting lack of co-localization with the nucleus.

**Supplemental Figure 5:**
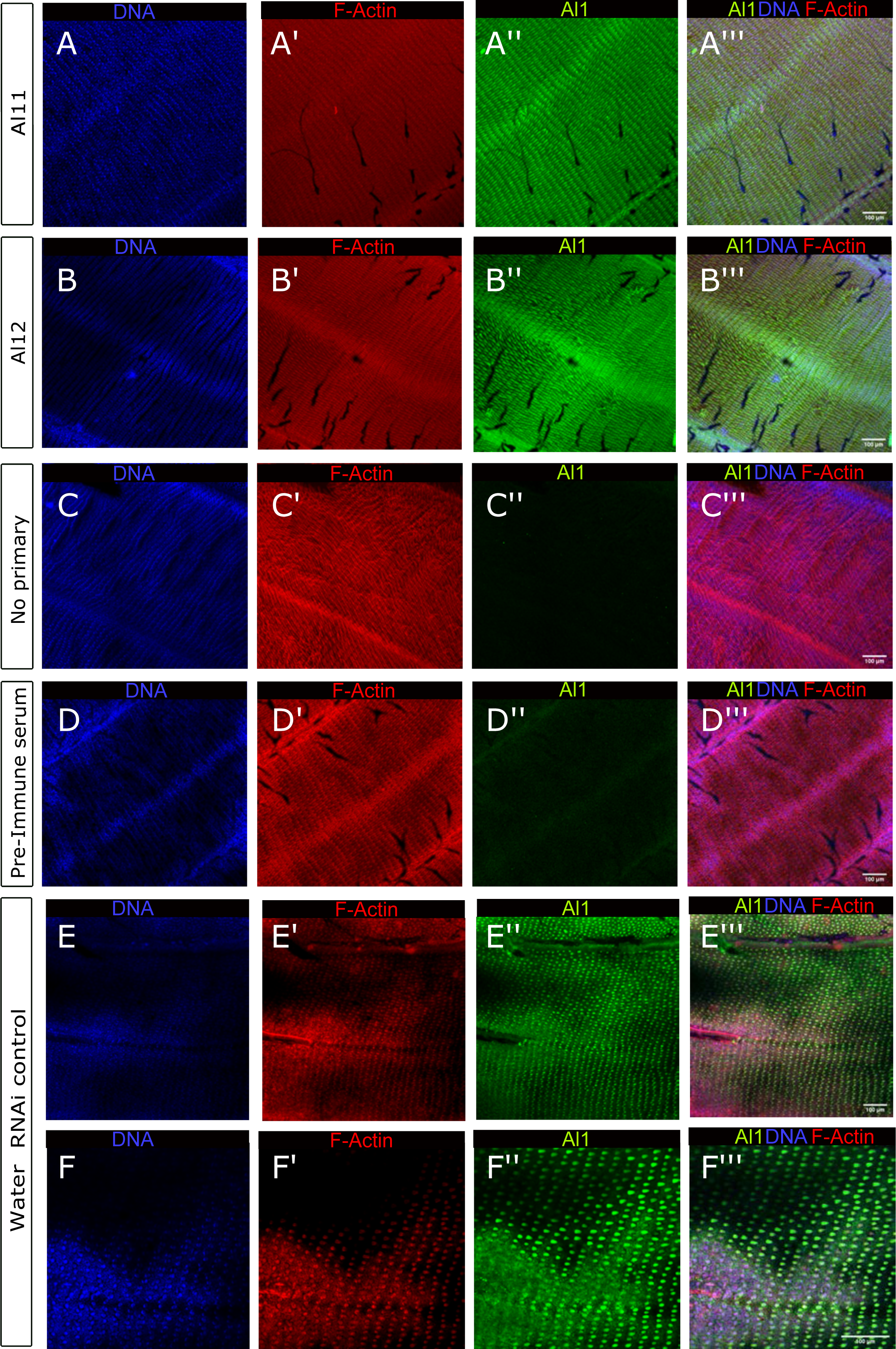
Immunodetection of Aristaless1 in white *Heliconius cydno* pupal forewings (between 2-3 days APF) across several control setups. (**A**) View of Al1 detection in scales by using the Al11 specific antibody (antibody used for all the immunodetections assays shown in the manuscript). (**B**) Al1 detection in scales by using the Al12, a different Al1 specific antibody targeting another part of the protein. (**C**) Negative control wing without any primary antibody. (**D**) Negative control wing in which the primary antibody was substituted by the pre-immune serum. (**E-F**) Al1 Immunodetection after a control water injection and electroporation. (**E**) View of an extended portion of the wing. (**F**) Closer view of scale cells to highlight details of Al1 protein detection following the manipulation. Panel show detection of DNA (A-F), F-actin (A’-F’), Al1 (A’’-F’’), and merge (A’’’-F’’’). The water injection site is located on the right corner outside of the field of view of the image.

**Supplemental Figure 6:**
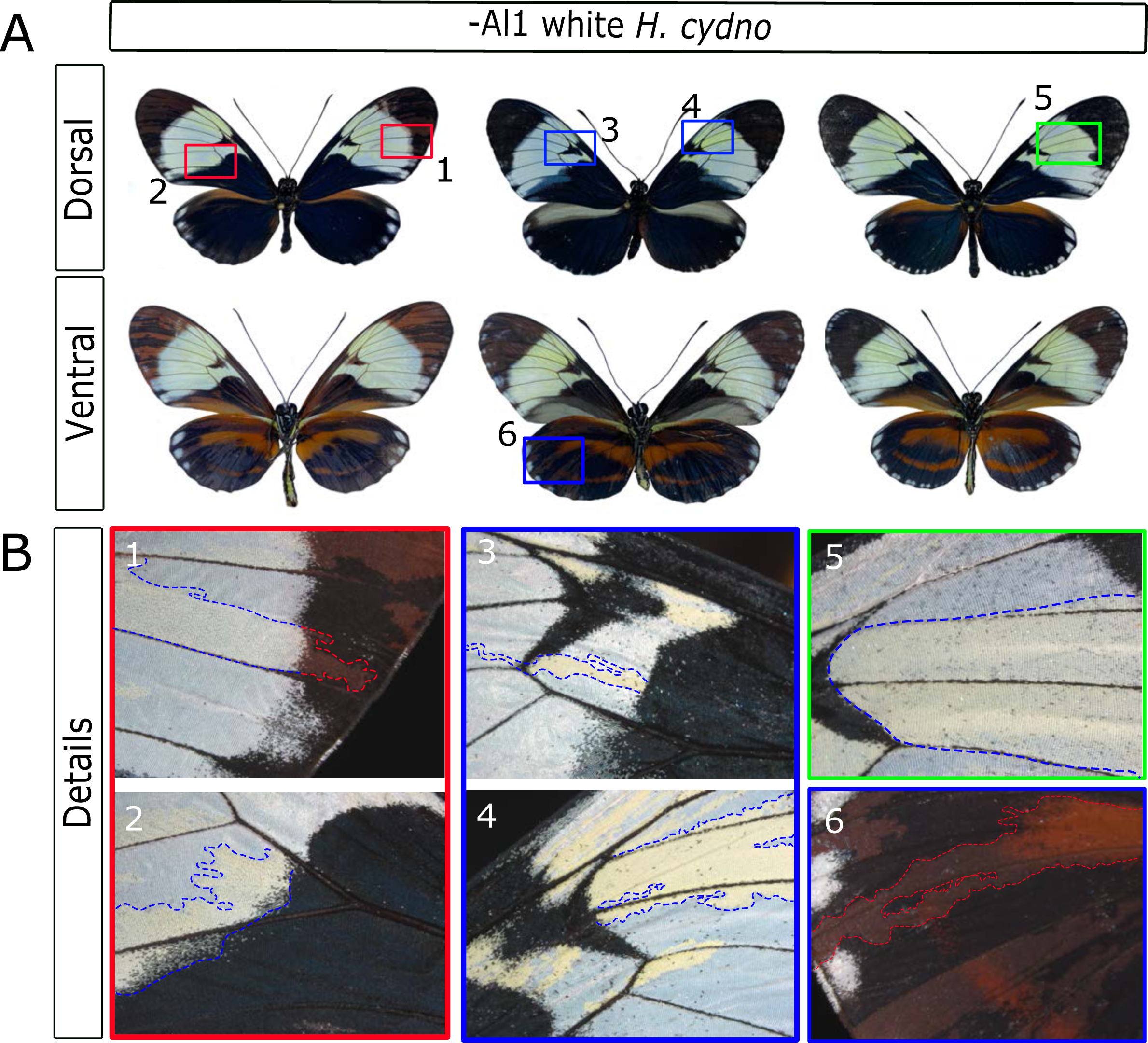
Showcase of the clones variation in Al1 CRISPR adults. **(A)** Whole butterfly views (both dorsal and ventral sides) of adults with Al1 CRISPR clones. The numbered squares are highlighted as closer views of the clones **(B)**. Some of the clones in which scales shift from white to yellow are highlighted by the blue dotted line and the clones in which scales shift from black to brown are highlighted by the red dotted line.

**Supplemental Figure 7:**
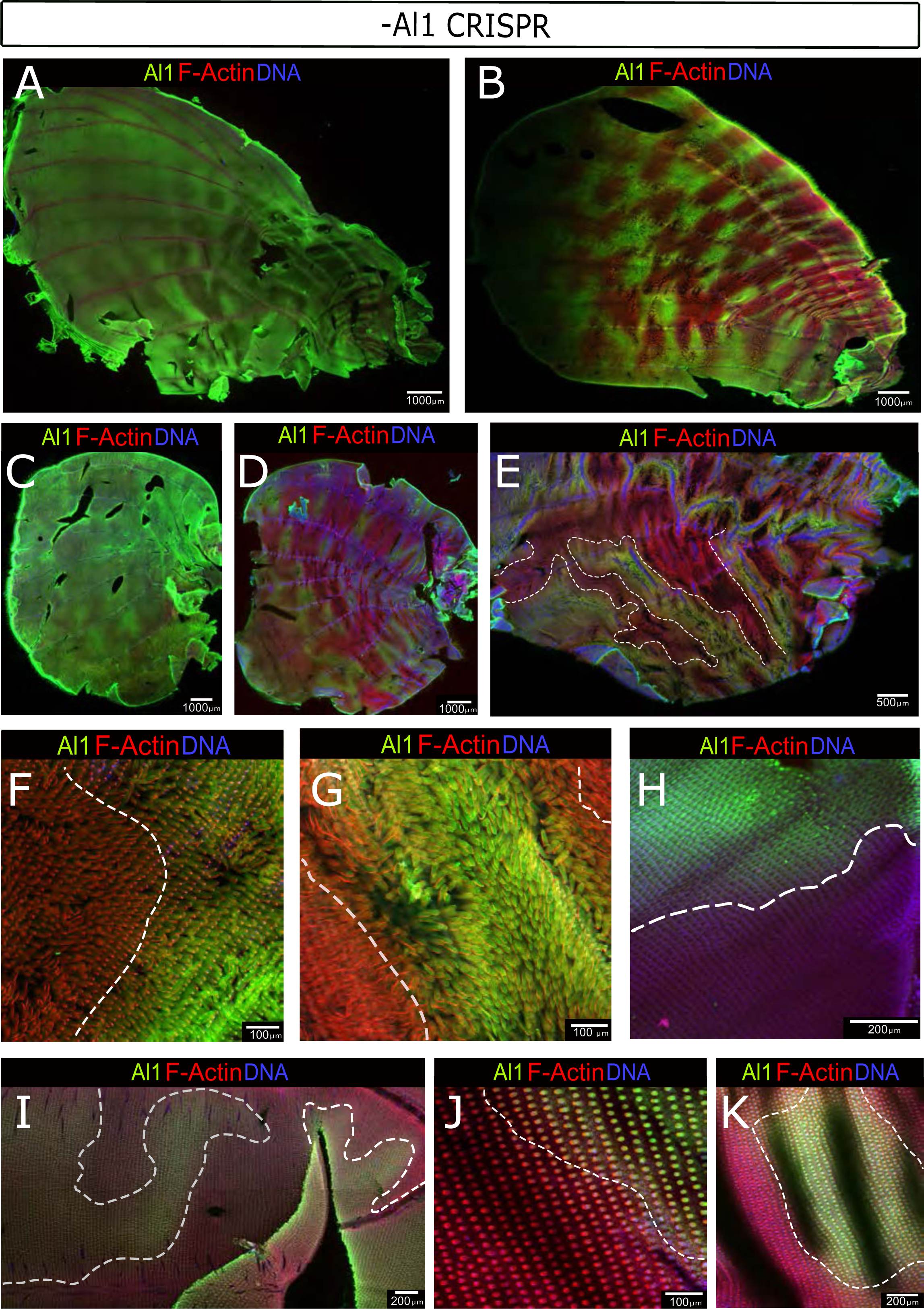
Showcase of the clones variation by immunodetection in Al1 CRISPR pupal wings (48 to 72 APF). (**A**) Low density to no clone forewing. (**B**) High clone density forewing highlighting scales lacking Al1. (**C**) Low density to no clone forewing. (**D**) High clone density hindwing highlighting scales lacking Al1. (**E**) Another High clone density forewing in which the clones have been highlighted with a white dotted line. (**I-K**) Details across multiple wings of different stages (48 to 72 APF) are shown to highlight the lack of Al1 within the clones. In all the detail views the boundaries are shown with a white dotted line.

**Supplemental Figure 8:**
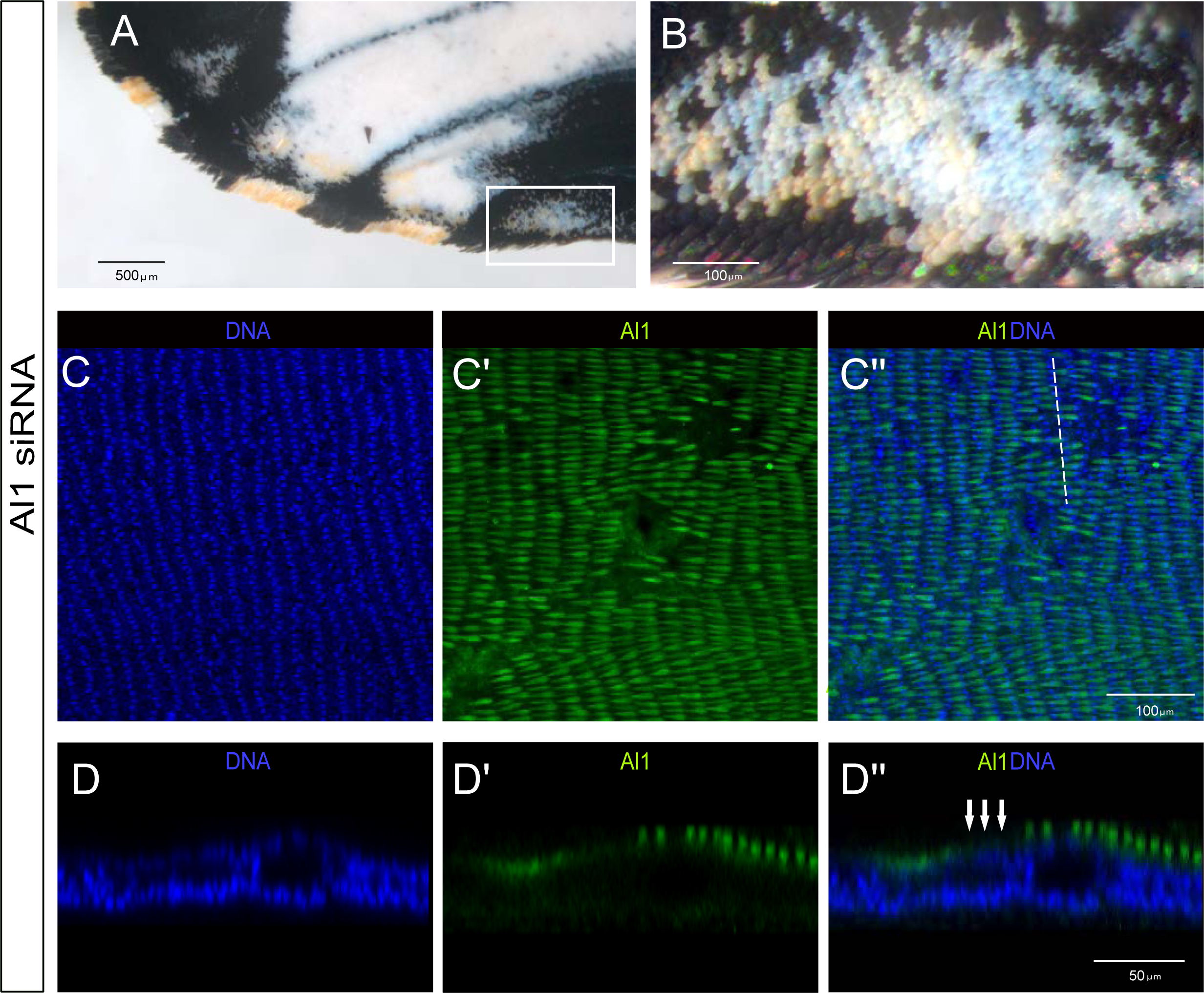
Immunodetection of Aristaless1 in *al1* RNAi knockdown pupal forewings of white *Heliconius cydno* (3 Days APF). (**A**) *al1* knockdown adult wings showing areas of the wing switching from white scales to yellow scales. (**B**) Higher magnification of the white square shown in A. (**C**) Immunodetection of Al1 in an *al1* knockdown pupal imaginal disc (3 days APF) showing patches of reduced or absent Al1 localization. (**D**) Side digital reconstruction (white dashed line indicate cross section in C’’) from a z-stack of one of the patches in panel C to observe scale morphology and the absence of *al1* in presumptive affected scales. Panel show detection of DNA (C,D), Al1 (C’,D’) and a merge (C’’, D’’) view. Scale bars: A, 500 μm; B-C, 100 μm; D, 50 μm.

**Supplemental Figure 9:**
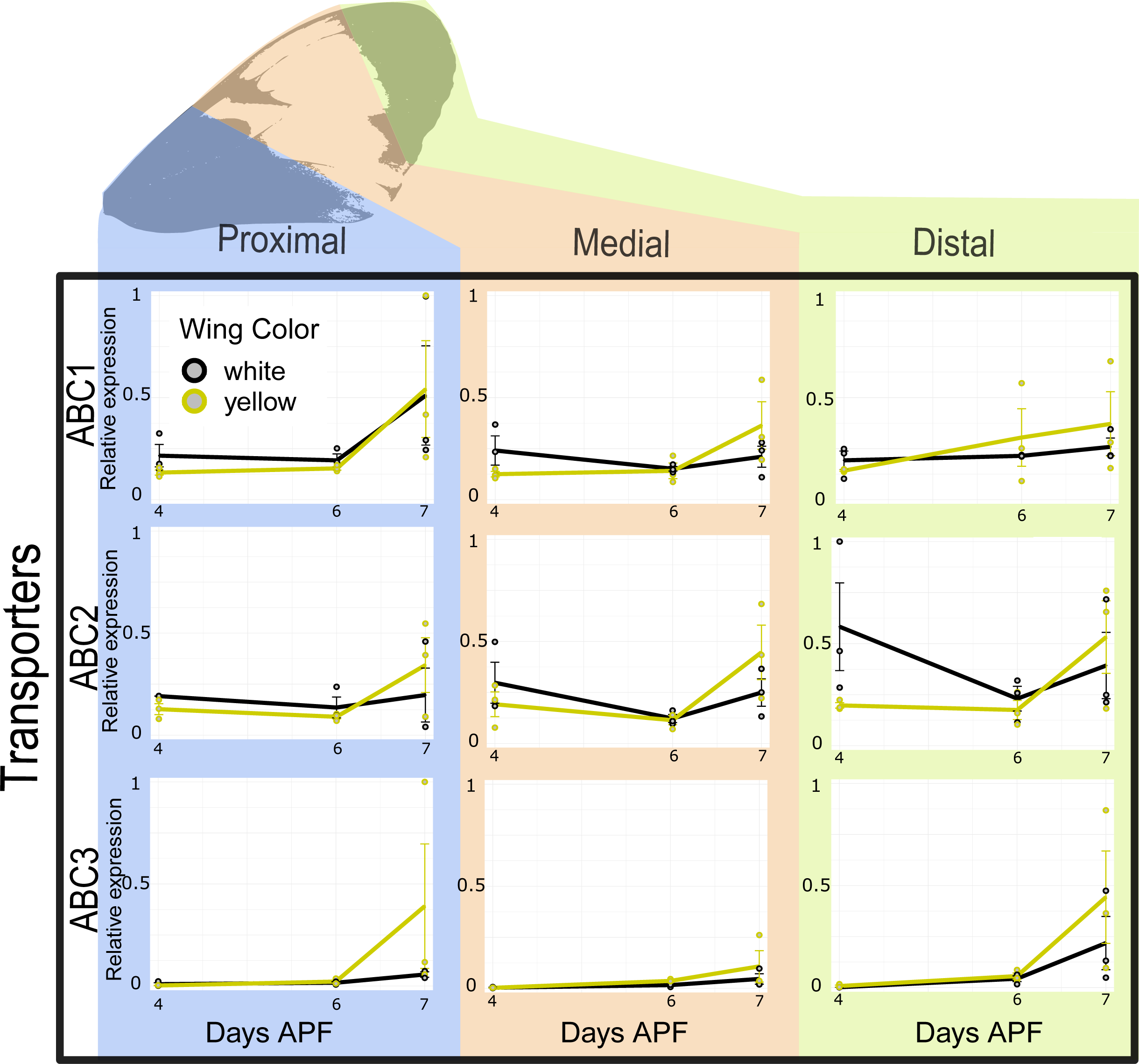
Downstream ABC transporters qPCR expression analysis between white and yellow *H. cydno* butterflies. Relative expression levels of each of the analyzed ABC transporters in white and yellow pupal forewings sections (proximal, media, distal) across 3 different time points (4, 6, 7 days APF). The relative expression values are scaled to the highest value across the wing sections for each one of the genes. The significance in the observed differences was tested using t-test. None of tested differences showed significance.

## Supplemental Table

**Table S1:**
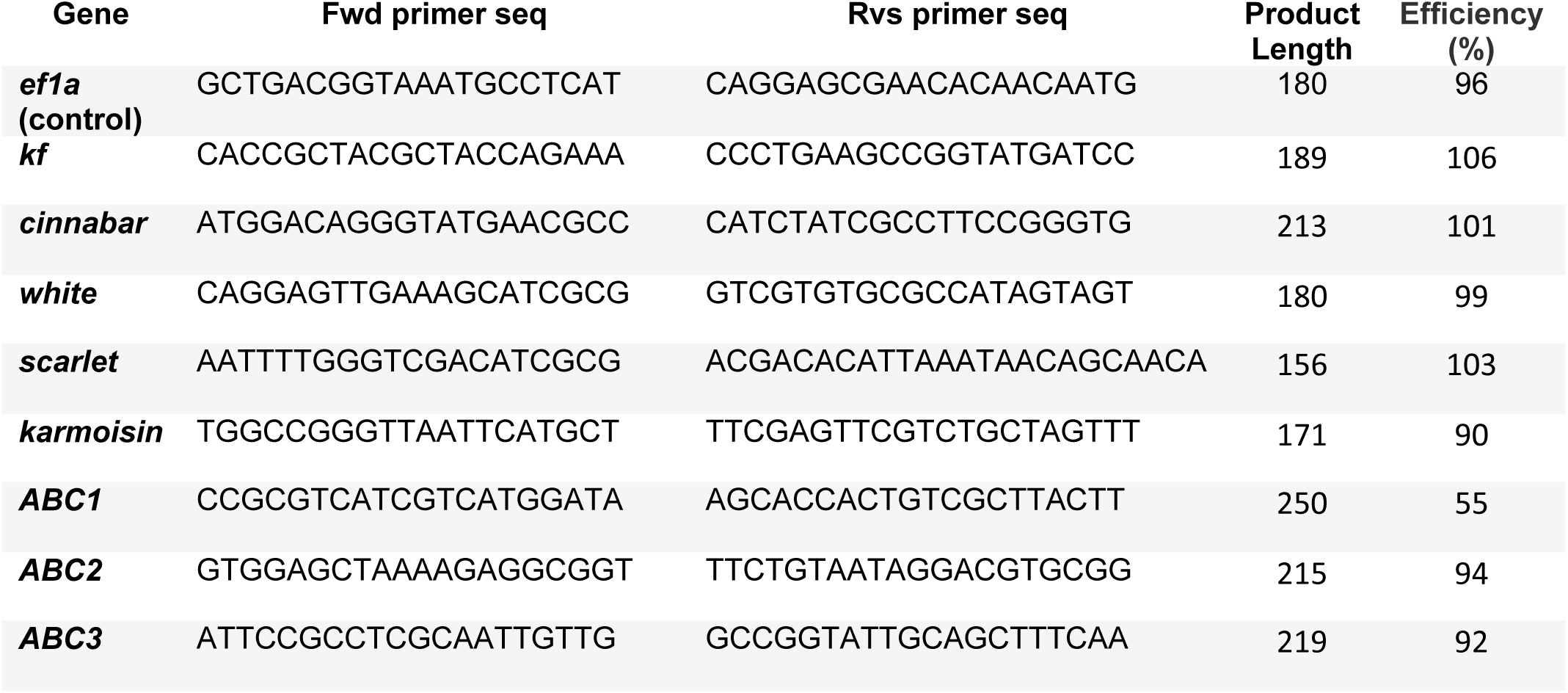
qPCR gene primers and efficiency tests.

## Source Data Files

**Source Data File 1: Zip file with the raw unedited blot images.** Sections of these 3 images were used to create Supplemental Figure 2.

**Source Data File 2: Uncropped Blot images with details on each section of the blot.** Sections of these 3 images as well as part of the data in the associated tables were used to create Supplemental Figure 2.

**Source Data File 3: Raw ΔCq data from our qPCR analysis used to calculate the relative expression of genes of interest.** The data of these source file is used to produce the plots in Figure 7 and Supplement Figure 9 ad described in the method section.

